# An efficient timer and sizer of biomacromolecular motions

**DOI:** 10.1101/384511

**Authors:** Justin Chan, Hong-Rui Lin, Kazuhiro Takemura, Kai-Chun Chang, Yuan-Yu Chang, Yasumasa Joti, Akio Kitao, Lee-Wei Yang

## Abstract

Life ticks as fast as how efficiently proteins perform their functional dynamics. Well-folded/structured biomacromolecules perform functions via large-scale intrinsic motions across multiple conformational states, which occur at timescales of nano-to milliseconds. Computationally expensive molecular dynamics (MD) simulation has been the only theoretical tool to gauge the time and sizes of these motions, though barely to their slowest ends. Here, we convert a computationally cheap elastic network model (ENM) into a molecular timer and sizer to gauge the slowest functional motions of proteins and ribosome. Quasi-harmonic analysis, fluctuation-profile matching (FPM) and the Wiener–Khintchine theorem (WKT) are used to define the “time-periods”, *t*, for anharmonic principal components (PCs) which are validated by NMR order parameters. The PCs with their respective “time-periods” are mapped to the eigenvalues (λ_ENM_) of the corresponding ENM modes. Thus, the power laws *t*(ns) = 86.9λ_ENM_^−1.9^ and σ^2^(Å^2^) = 46.1λ_ENM_^−2.5^ are established allowing the characterization of the time scales of NMR-resolved conformers, crystallographic anisotropic displacement parameters, and important ribosomal motions, as well as motional sizes of the latter.

**GRAPHICAL ABSTRACT:** 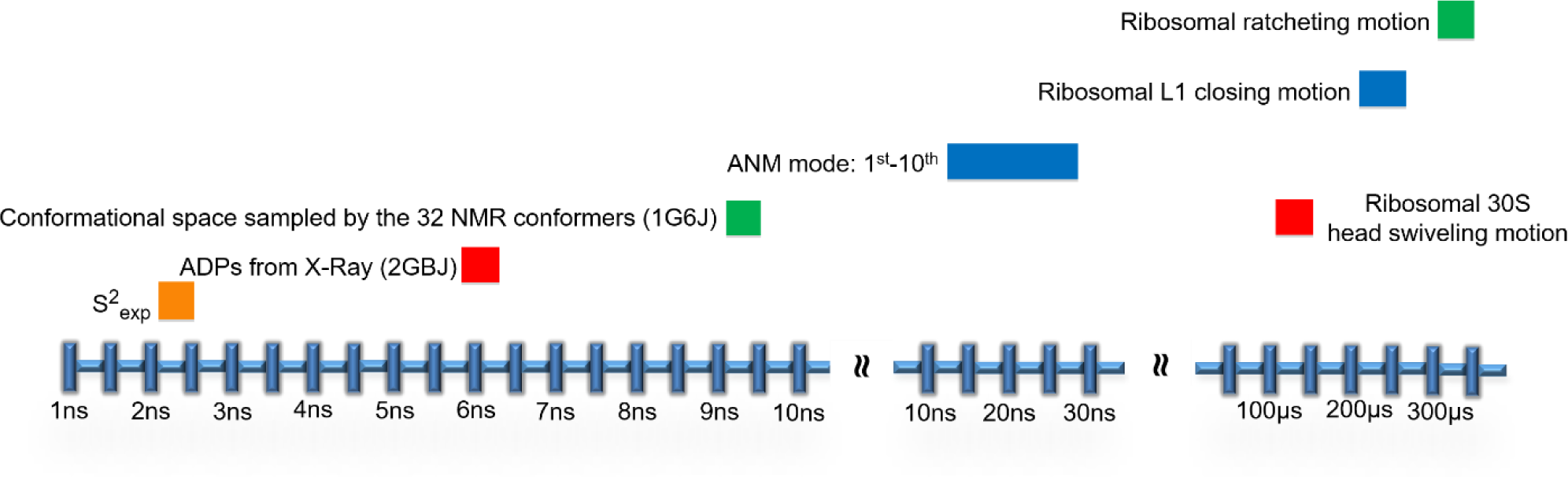

## INTRODUCTION

The magnitude of protein functional motions for folded proteins (or ‘equilibrated’ proteins) modulate the rates of their underlying physiological processes (Popovych et al., 2006). In principle, the wider the conformational spread is, the longer the time it takes, although the time and space relationship for protein motions damped in water may not be readily described by free diffusion of harmonic oscillators (Atilgan et al., 2001; Bahar et al., 1998; Flory et al., 1976; Li et al., 2016; Tirion, 1996; Wang et al., 2004). Understanding such a relation can bring important insights in protein dynamics including but not limited to the time scales of molecular controls as well as sizes of conformational changes that support the molecular function or facilitate two-body interactions. Molecular dynamics (MD) simulations, though computational costly, has been the only option to explore such a relationship (Götz et al., 2012; Ratje et al., 2010; Sanbonmatsu et al., 2005; Shaw et al., 2010; Whitford et al., 2010), while larger and time-wise lengthier motions (>10s of microseconds for large macromolecules) are still not within the reach of modern MD.

On the other hand, computationally cheap (>5 order cheaper than MD) elastic network models (ENMs) have been widely used for more than two decades (Atilgan et al., 2001; Bahar et al., 1998; Flory et al., 1976; Tirion, 1996) to study and predict the span (variance) of spatial distributions of protein conformations accessible to a molecule. These physically intuitive models have been used to study vibrational dynamics of all the proteins/supramolecules (Li et al., 2016; Wang et al., 2004) in Protein Data Bank (PDB) where the “directions” of proteins’ conformational changes as observed by x-ray crystallography have been satisfactorily reproduced in hundreds of applications and in database-wide studies (Bahar et al., 2010; Li et al., 2017) (see the caption in **Figure 1**). However, the absolute time scales and variances of functional modes cannot be directly assessed by ENM due to its harmonic approximation. Functional modes, particularly, involve motions at large scales, wherein proteins traverse across multiple intermediate states (corresponding to local traps in the energy landscape; see the left and middle in **Figure 1**), and therefore are anharmonic in nature (Kitao and Go, 1999; Kitao et al., 1991; Yang and Chng, 2008). In other words, the true challenge lies on how to properly define the “time periods” of these anharmonic modes, which cannot be simply inferred from the length of the simulation (a small protein can travel several times on a given normal mode in a long MD simulation). Once defined, efficient and accurate methods are needed to predict the “*time periods*” and *absolute variances* of these anharmonic modes.

**Figure 1.**
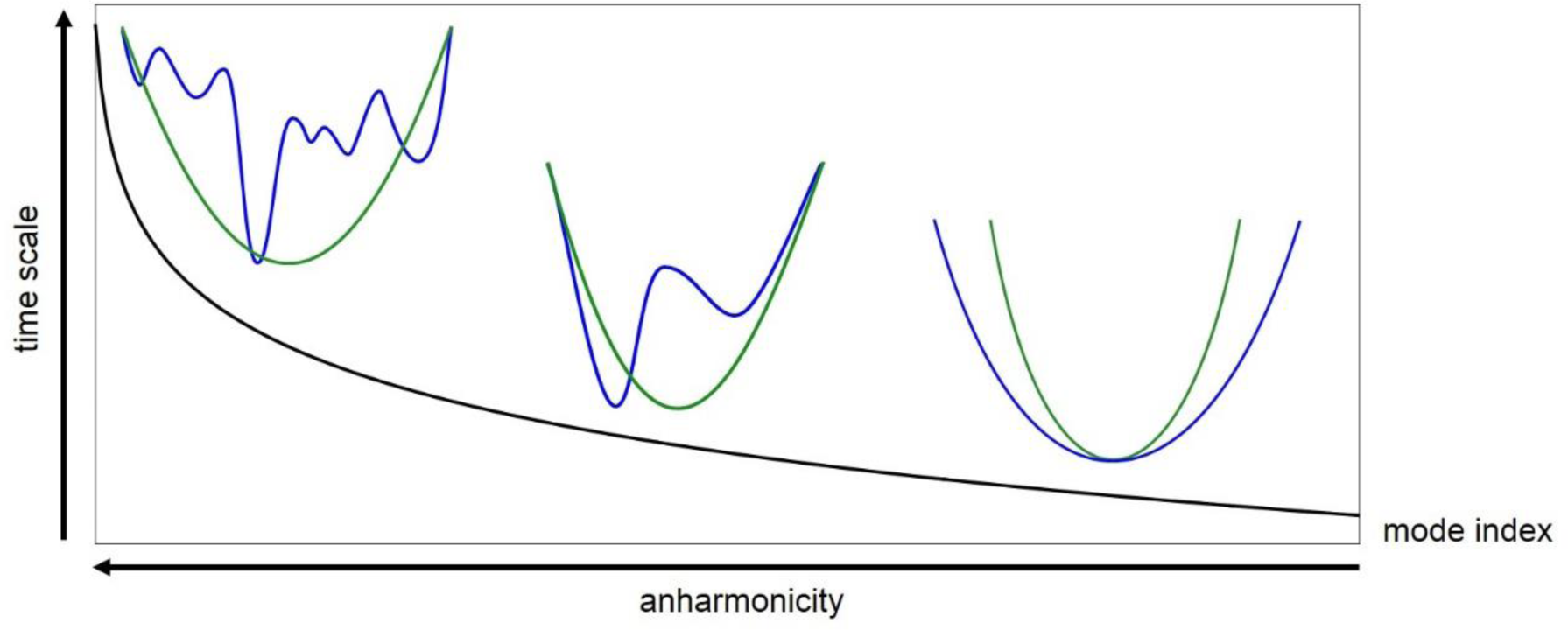
Time scales versus anharmonicity of the modes. The blue and green curves indicate the real and harmonically approximated energy landscapes, respectively. The slower the modes are, the larger the deviations from a simple harmonic approximation of the energy landscape (equivalently stated as having enriched anharmonicity (Kitao and Go, 1999; Kitao et al., 1991; Yang and Chng, 2008)). ENM-approximated harmonic energy landscape forms an “envelope” that outlines the real anharmonic energy landscape (Li et al., 2017) (to the left in **Figure 1**), which explains the observed correspondence between the theoretically predicted and experimentally characterized “directions” of conformational changes. However, the time scales and absolute spatial span of the modes, among the slowest and often functional, cannot be satisfactorily assessed given the harmonic approximation, which motivated us to design the current method to address the issue. The time scale estimated using the Intensity Weighted Period (IWP) method is introduced in this work (see below).

In this study, we first use Principal Component Analysis (PCA) of MD trajectories, time-correlation functions, Wiener–Khintchine theorem (WKT) (McQuarrie, 2000) to define “time periods” for all the anharmonic and harmonic modes. Then, we use a mapping technique, Fluctuation Profile Matching (FPM), to map a dynamic variable (containing information of spatial spread of protein atoms; see below) or an ENM mode to a Principal Component (PC); hence the time scale of that PC can be assigned to the dynamic variable or ENM mode of interest. When every ENM mode is assigned a time scale or MD-sampled conformational spread (variance), we can then derive corresponding power laws to describe how the time scale and conformational spread of a normal mode can be functions of its eigenvalue.

## RESULTS

### Overview

As a methodological overview in finer detail, **Figure 2** describes the autocorrelation function of a PC mode of ubiquitin. As previously described in either position (Okan et al., 2009) or velocity (Kitao et al., 1991) autocorrelation functions, the function decays exponentially as a damped harmonic oscillator (**Figure 2**). We tried three methods to define the period, or “timescale” in this context, of this damped harmonic oscillator for a PC mode. Among them, the Intensity Weighted Period (IWP; see definition below) requires the least assumption and appears to best describe the observed oscillatory behavior (**Figure 2**). Next, FPM identifies subsets of PCs that best reproduce (per the highest correlation) the experimentally observed profile of a given dynamical variable (including root-mean-square-fluctuations (RMSF) of a structural ensemble (Yang et al., 2009), anisotropic displacement parameters (ADPs) (Eyal et al., 2007) and NMR order parameters (Tjandra et al., 1995)) and assign the IWP (or “time scale”) of the slowest PC as the timescale of that profile. The subsets of PCs are generated by removing the top-*k* slowest PCs derived from long MD simulations. This is because we have previously (Yang et al., 2007) shown that deletion of the slowest ENM modes does not change the correlation between the predicted and experimentally observed B-factors, suggesting that dynamical profiles such as B-factors or anisotropic displacement parameters (ADP) do not include the protein’s slowest motions given the confined crystalline environment (Yang et al., 2007). In this study, we employ one of the most widely used ENMs, the anisotropic network model (ANM) (Atilgan et al., 2001; Chandrasekaran et al., 2016; Li et al., 2014). Note that it was known that the mapping of PCA mode and ANM mode (or other ENM mode) is not a one-to-one relationship; a slow PC (ANM) mode is best interpreted by a few slowest ANM (PC) mode (Bakan and Bahar, 2009; Nicolay and Sanejouand, 2006). FPM provides a robust mapping between a PC mode *k* and an ANM mode *k’* (or an experimental variable) when the variance comprising all the PC modes higher than *k* has the highest correlation with the variance comprising all the ANM modes higher than *k’* (see **STAR Methods** for FPM’s mathematical definition). The experimental variables here include NMR order parameters, ADPs from the X-ray structure and RMSF of NMR-solved structural ensemble of ubiquitin. To use the ANM as a convenient molecular timer and sizer, we match each ANM mode to a PC mode using FPM. Then, the timescale and variance of an ANM mode are assigned by the timescale and variance of the matched PC mode. We observe that the eigenvalues of ANM modes has a power law relationship with their assigned timescales and variances. Applying the aforementioned procedure to three protein systems, ubiquitin, FGF2 and HP-NAP, we derive master equations for time and size of protein motions as power laws of ANM eigenvalues. We confirm the size power law by reproducing the observed ribosomal body rotation and estimate the time-scales for ribosomal body rotation and head swiveling motions to a narrower time range than those suggested by MD simulations and experiments.

**Figure 2.**
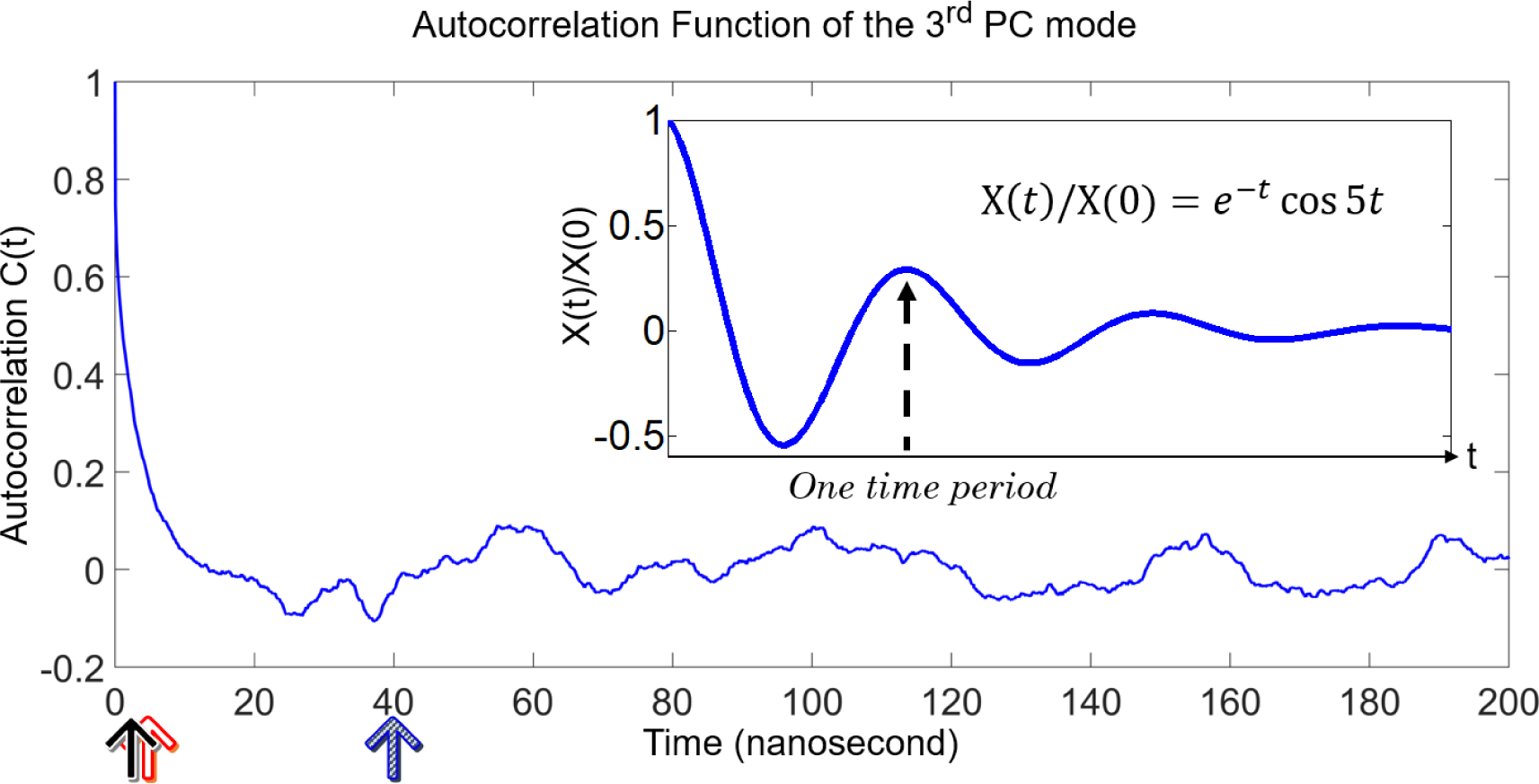
Defining the time scale of a PC mode using three methods. The profile of the autocorrelation function for the third PC mode, derived from PCA of a 600-ns simulation of ubiquitin (PDB ID: 1UBQ) and Wiener–Khintchine theorem (**Equation 1**), is plotted. Three time constants *τ*_*c*_ = 2.34 ns (indicated by the black solid arrow), *τ*_*r*_ = 2.79 ns (red hollow arrow; obtained using fitting in the time domain) and *τ*_*w*_ = 40.45 ns (blue shaded arrow) that characterize the exponential decay of the autocorrelation function *C*(*t*) = *A* exp(–*t*/τ_*r*_), characteristic time such that 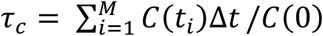 and IWP of the power spectrum that is 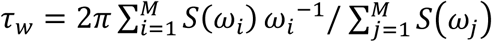, respectively, are used to describe the vibrational relaxation process that mimics a damped harmonic oscillator. In addition, because the Fourier transform of an exponential function is a Lorentzian function (Boas, 2006), we can also obtain *τ*_*r*_ from fitting the entire power spectrum using a Lorentzian function. However, the resulting *τ*_*r*_ is much smaller than *τ*_*c*_ with a fitting correlation close to zero (data not shown). **(Inset)** Theoretical profile of one particle oscillating in an underdamped regime. The equation comparatively shows the equivalent decay and oscillation rate of the autocorrelation function. The vertical arrow indicates the time period, which in this illustrative example is 2π/5.

### Determining the time scales of the anharmonic modes

To estimate the true time scale of a PC mode, we first performed a 600-ns MD simulation for the 76 amino acid signaling protein, ubiquitin (Vijay-kumar et al., 1987) (PDB ID: 1UBQ). PCA analysis (Yang et al., 2009) was carried out on the trajectory of the first 72 residues of ubiquitin, which are structurally ordered to derive the covariance matrix <Δ**R**Δ**R**^***T***^> and the projection of each snapshot to each of the eigenvectors (see details in the **STAR Methods**). In other words, a snapshot is a scalar value (PC mode coordinate) on the mode *k.*, **V**_*k*_. The total *M* snapshots projected on mode *k* together with padded *M* zeros result in a projection series *u*_*k*_(*s*) = {*u*_*k*0_, *u*_*k*1_, … *u*_*kM*–1_, 0^*M*^, 0^*M*+1^ … 0^2*M*–1^}, where *u*_*kj*_ means the *j*-th snapshot projected on the mode *k*, based on which the autocorrelation function for the mode *k* is calculated as follows. Let 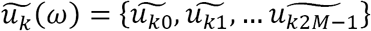 be the Fourier transform of *u*_*k*_(*s*), and the spectral density of the process *u*_*k*_(*s*) can be defined as 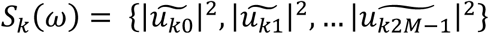. According to the Wiener–Khintchine theorem (McQuarrie, 2000), the autocorrelation function of *u*_*k*_(*s*), *C*_*k*_(*t*), can be calculated as the inverse Fourier transform of the spectral density *S*_*k*_(ω) such that

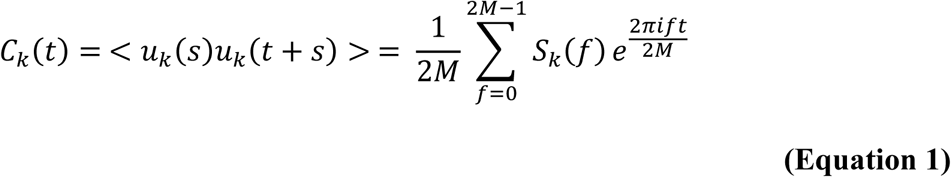

where, *f* = ω/2*π* is the frequency. The mathematical details and computational implementation are provided in the **STAR Methods**.

**Figure 2** shows that the *C*_3_(*t*) of the third PC mode, where *C*_3_(0) is unity (after normalization) at time zero, drops to where there are negative correlations and rises back above zero before eventually vanishing in an oscillatory manner. Most of the slowest modes demonstrate such a behavior, which can be approximated as a harmonic oscillator in underdamped regimes (illustrated in the inset of **Figure 2**). Our purpose here is to define a reasonable “time period” for such a process, which will be inferred as *the time scale (or “one period”) of mode k*.

We initiated three attempts to define the time period of the PC mode. For the first one, we approximated the relaxation of *C*(*t*) from unity to zero (the first time it hits a value barely larger than zero) as an exponential decay with a relaxation time constant of *τ*_*r*_. Because the Fourier transform of an exponential function is a Lorentzian function (Boas, 2006), we can also obtain *τ*_*r*_ from fitting the entire power spectrum using a Lorentzian function, in contrast with fitting *C*(*t*) within a limited time range [when *C*(*t*) > 0]. For the second method, we define the characteristic time constant *τ*_*c*_, as the integration of *C*(*t*) over the simulation time divided by *C*(0) which is equal to *τ*_*r*_ if *C*(*t*) is indeed an exponentially decaying function. For the third attempt, we simply take the intensity-weighted average of the periods in the power spectrum of *S*(ω) such that:

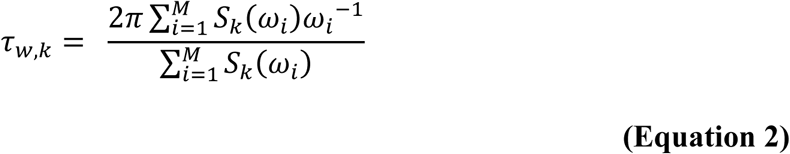

where *S*_*k*_(ω_*i*_) is the intensity (or “weight”) for the frequency component *ω*_*i*_ (*ω*_*i*_^−1^ is its corresponding period). As shown in **Figure 2**, *τ*_*w*_, the intensity-weighted period (IWP), reaches a time scale where the *C*(*t*) is close to the peak of the first wave, resembling a time period of a damped oscillator, while neither relaxation time *τ_r_* (obtained from the fitting in both time and frequency domains) nor characteristic time *τ_c_* covers the time span for a period of an approximated damped oscillator.

### Determining the time scales of experimentally observed dynamical variables

The spatial distribution of every residue (or every atom) in a protein near its folded (equilibrated) state can be observed by Cα RMSF of a NMR-resolved structural ensemble (Yang et al., 2009), NMR order parameters (Tjandra et al., 1995) and x-ray crystallographical anisotropic displacement parameters (ADPs) (Eyal et al., 2007). Plotting the dynamical value for each residue against its residue index results in so-called “observed fluctuation profiles” (oFPs) (**Figure 3, top**, black circles). Every residue has one magnitude of RMSF, one order parameter and 6 ADP values (including variance xx-, yy-, zz- and covariance xy-, yz- and xz-components (Eyal et al., 2007)). Concurrently, we derive the theoretical counterparts of these measured variables from a MD snapshot-derived covariance matrix, <Δ**R**Δ**R**^*T*^>_*MD,k*,_ comprising all the PC modes ≥ *k*^*th*^ mode (lower PC modes have larger variance)

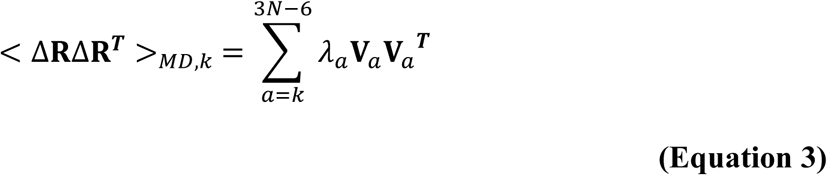

where **V**_*a*_ is the eigenvector with its corresponding eigenvalue, λ_*a*_, taken from the diagonal of **Λ** in **Equation S1** in the **STAR Methods** for the *a*’th PC mode; *N* is the total number of heavy atoms in the protein.

**Figure 3.**
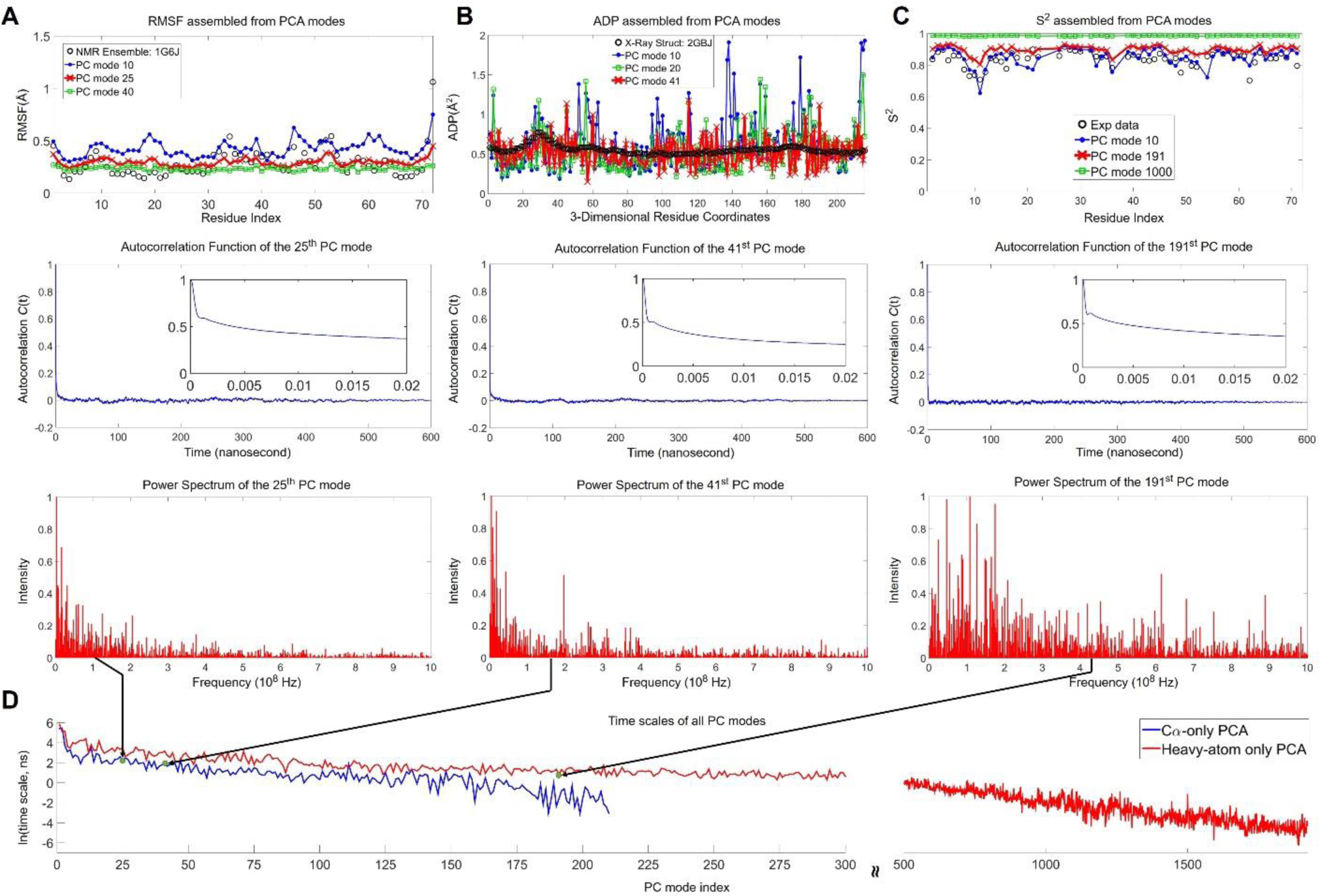
Time scale mapping for different experimental data that characterize protein dynamics. The first rows for **(A-C)** are, respectively, the oFP of RMSF derived from an NMR-determined ensemble (PDB: 1G6J), ADP values (only the diagonal elements are shown) and NMR-measured order parameters against residue index of ubiquitin, where the experimental profiles are plotted in scattered black circles, and the best fit PC mode profile is plotted in thick red lines. **(A, top)** The experimental profile best matches (σ_p_ = 0.69) the theoretical RMSF derived from a 600-ns MD simulation, where all PC modes ≥25 are included to contribute to <Δ**R**Δ**R**^***T***^>_*MD,k*_ (*k* = 25); further removing modes higher than mode 24 would impair the correlation. Profiles of different *k* values are also plotted for comparison. **(A, middle)** The autocorrelation function for the best-matched mode *k* = 25 is computed via WKT following **Equation 1**. In addition, **(A, bottom)** its corresponding power spectrum, *S*(ω), plotted against its constituent frequencies *f* = ω/2*π*, can be obtained before the inverse Fourier transform. Only the contributions of frequencies lower than 10^9^ Hz are plotted. A black arrow is drawn to point from the implied IWP’s frequency, (1/*τ*_*w*_), to its corresponding IWP (*τ*_*w*_) in the ordinate (taken natural logarithm) of the chart in panel **(D)**. **(B and C)** In a similar spirit as **(A)**, panels **(B and C)** are plotted for the experimental ADPs (PDB: 2GBJ) and NMR order parameters (Tjandra et al., 1995) of ubiquitin, with the best matched modes determined as *k*=41 and *k*=191, respectively. Note that the PCAs performed for **(A and B)** are Cα-based and for **(c)**, the PCA is based on all-heavy-atoms and H atoms bonded to backbone N’s. **(D)** IWP (*τ*_*w*_) is plotted against the PC mode index for both Cα-based (3*N* – 6 = 210 modes; in blue) and all-heavy-atom based (3*N* – 6 = 1923 modes; in red) PCA.

Next, we use FPM to find the *k* that can best reproduce the oFPs of interest, we seek the optimal k where the profile derived from <Δ**R**Δ**R**^*T*^>_*MD,k*_ has the best agreement with the oFPs by having the highest Pearson’s correlation coefficient (σ_p_) (Pearson, 1895). After finding the best *k* via fluctuation profile matching (FPM), *τ*_*w,k*_ of the PC mode *k* is assigned to the observed dynamical variable, as its estimated time scale (see **Figure 3** and **Supplemental Information** for examples).

The NMR order parameters, *S*^2^, describes the order of the backbone -NH bond vector *r*_*ij*_ = (*x*_*ij*_, *y*_*ij*_, *z*_*ij*_) pointing from atom *i* to *j* (herein, *i* is N, and *j* is H) which can be approximated as (Best and Vendruscolo, 2004; Henry and Szabo, 1985; Yang et al., 2009)

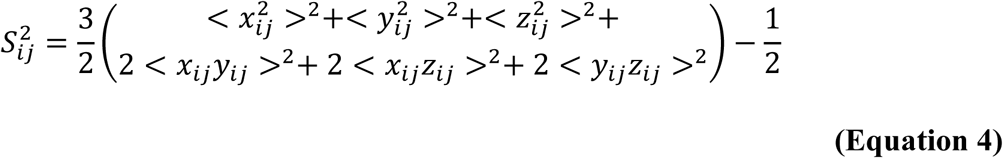

Here, the length of *r*_*ij*_ is normalized to unity. The angular brackets denote averages taken over the *M* snapshots; herein *M* = 6,000,000 or 1,200,000 are for the 600 ns and 120 ns trajectories, respectively. *S*^2^ takes on values between 0, for freely rotated bonds (disordered), and unity, for perfectly “ordered” bonds. PCA is carried out for all the heavy atoms plus the hydrogen atoms in the amino (–NH) groups along the backbone of ubiquitin on the long 600-ns MD simulations and the associated five-constituent 120-ns sub-trajectories.

Through FPM, it was found that *τ*_*w,k=*191_ = 2.34 ns and σ^2^_*k=*191_ = 0.44 Å^2^ (**Figure 3C** and see correlation as a function of PC mode index, peaked at the mode 191, in **Figure S2C**), characterizes the time scale and size of motion for the observed order parameter profiles (Tjandra et al., 1995). When we applied the current method to shorter trajectories, it was found among five 120-ns simulations that the *S*^*2*^_*exp*_ profile consistently maps to the PC modes with *τ*_*w*_ values between 0.7 ns and 1.0 ns (see one 120-ns result in **Figure S3A**).

Hence, despite different simulation lengths being used, our method consistently reports a time scale of 0.7 ns and 2.3 ns for this set of dynamics variables characterized by the NMR relaxation experiments, consistent with the correlation time of methyl symmetry axis motion (300 to 500 ps) (Lee et al., 1999) using the extended model-free approach, while the “back-calculated” analysis of spectral densities using MD simulations reported time constants of 0.6, 0.9, and 1.5 ns (Nederveen and Bonvin, 2005).

### Determination of time scales of the ENM modes and inference of time scales from the ENM eigenvalues

It is cumbersome to repetitively perform long MD simulations every time to characterize the time scales of experimentally observed dynamics variables. Here, we devised a computationally light molecular timer that can estimate the time scales of modes from anisotropic network model (ANM) (Atilgan et al., 2001; Li et al., 2017). To realize this, FPM of RMSF profiles derived from ANM modes and from MD PC modes was used to map each ANM mode to a PC mode. The ANM derived covariance matrix, <Δ**R**Δ**R**^*T*^>_*ANM,l*_, for all ANM modes ≥ *l* (lower ANM modes have lower frequencies or smaller eigenvalues) was calculated as follows

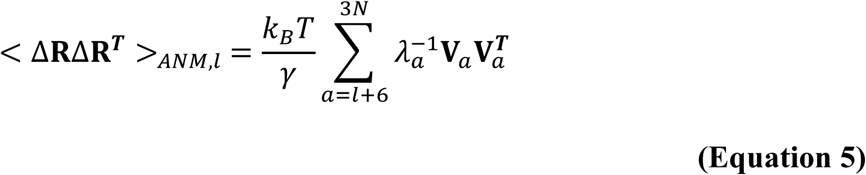

where ***V***_*a*_ and *λ*_*a*_ are the eigenvector and eigenvalue of the *a*’th ANM mode, respectively. While, γ is the universal spring constant, k_B_ is the Boltzmann constant and T is the absolute temperature with 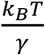 set to unity. The RMSF_*ANM,l*_ and ADP_*ANM,l*_ profiles were computed from the <Δ**R**Δ**R**^***T***^>_*ANM,l*_ matrix with the same methods used for MD-derived profiles of RMSF and ADP.

FPM is applied to each ANM mode *l*’s RMSF_*ANM,l*_ profile to identify the *k*^*th*^ PC’s RMSF_*MD,k*_ profile with the highest correlation. By doing this, every ANM mode *l* can be assigned the time scale (*τ*_*w*_) of its best-matched *k*^*th*^ PC mode. If there are more than one ANM mode mapped to a PC mode, the ANM mode having the highest correlation with that PC mode would be kept. We then fit the remaining ANM eigenvalues (*λ*_*ANM*_) with their mapped time scales (*t*_*ANM*_) in the form of a power law *t*_*ANM*_ = *c* × *λ*_*ANM*_^*d*^ to obtain the constants *c* and *d*, as described in the **STAR Methods** and **Figure 4A**.

**Figure 4.**
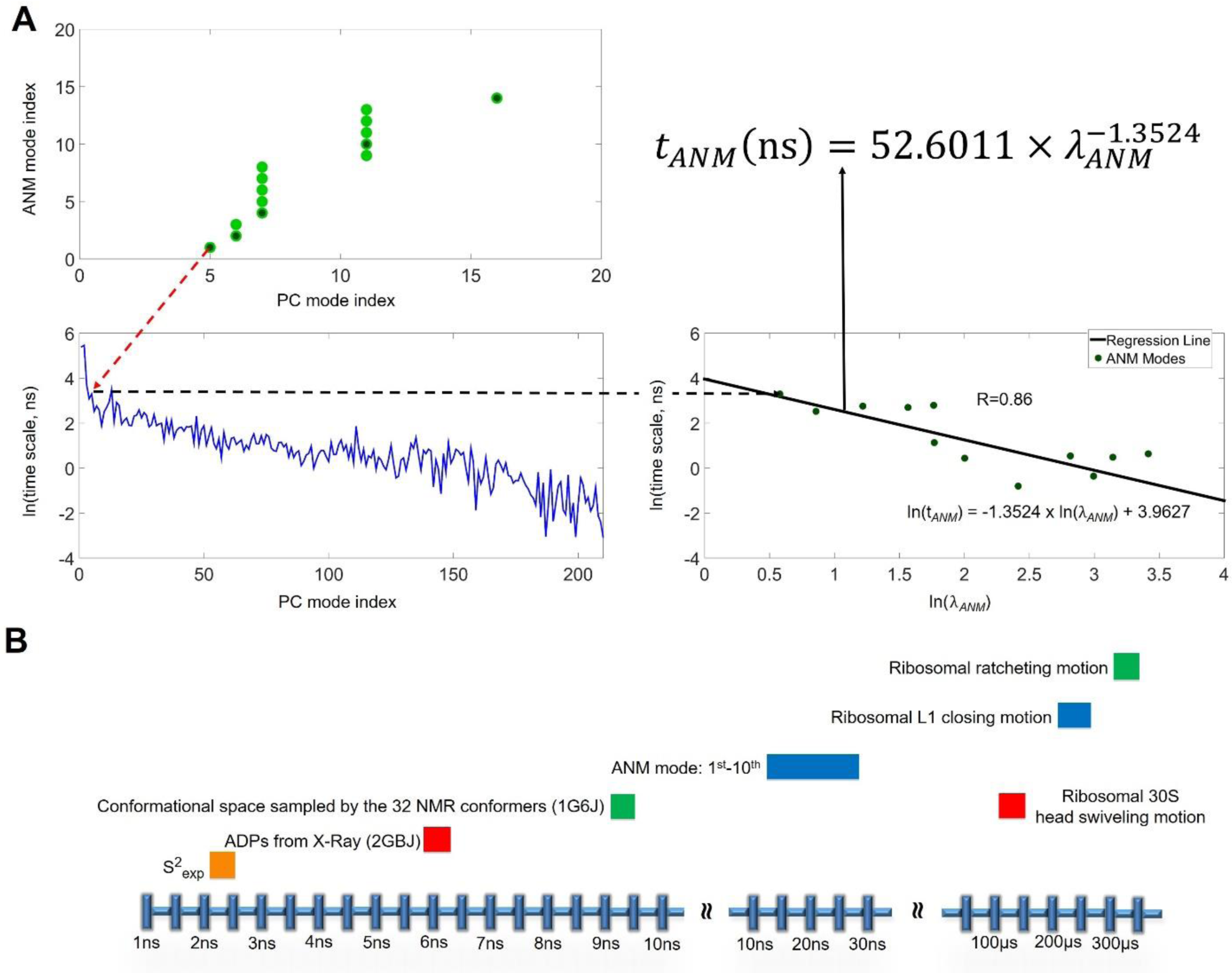
Frequency mapping for ANM to obtain the power law of *τ*_*w*_ as a function of ANM eigenvalues. **(A)** The ANM analysis and PCA of long MD trajectories are obtained for ubiquitin (residues 1-72, Cα-only). The *τ*_*w*_ of every PC mode obtained using IWP was mapped to the most correlated ANM mode using FPM. **(A, top left)**, First *l*-1 ANM modes are removed to obtain a covariance matrix comprising modes ≥*l*. The RMSF of each residue, RMSF_*ANM,l*_, is the square root of the sum of x-, y- and z-components of its variance. This RMSF_*ANM,l*_ is compared to the RMSF_*MD,k*_ comprising PC modes ≥*k*. Each ANM mode *l* (light-green circles) is mapped to the highest correlated PC mode *k* by comparing RMSF profiles. When multiple ANM modes are mapped to the same PC mode *k*, only the pair with the highest correlation is kept (dark-green circles). The *τ*_*w*_ of the PC mode *k* is then assigned to the ANM mode *l*. For example, the first (*l =* 1) ANM mode maps to the 5^th^ PC mode (red arrow). **(A, bottom left)**, The *τ*_*w*_’s are plotted against the PC mode index (identical to the blue curve in **Figure 3D**). **(A, bottom right)**, The *τ*_*w*_’s are plotted against λ_*ANM*_ (the eigenvalues of ANM modes). Linear regression gives the power law 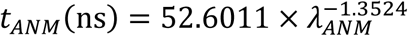 for ubiquitin. **(B)** Time scales of the examined dynamics variables, including the ANM modes, are assigned by the PCA+WKT+FPM approach for ubiquitin, except for the time scales of ribosomal motions that are estimated by the power law 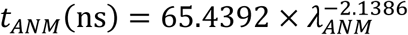 derived from the combined *τ*_*w*_ and λ_*ANM*_ of ubiquitin and FGF2.

For the case of ubiquitin, ANM mode 1 is mapped to PC mode 5 therefore it is assigned the time scale, *τ*_*w,k=*5_ = 27.03 ns (**Figure 4A**). Similarly, the RMSF_*ANM*_ for modes 4 and 10 are mapped to PC modes 7 and 11 with the corresponding time periods *τ*_*w*_ of 21.95 and 14.87 ns, respectively. This suggests that the motions described by the first 10 ANM modes for ubiquitin, or the slowest end of ANM, should occur within the timeframe of 13 to 28 ns (see **Figure 4B**). Fitting the *λ*_*ANM*_’s with the *t*_*ANM*_’s we obtain a power law for ubiquitin,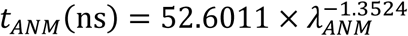.

To extend the applicability of this method, we apply the aforementioned method to two other proteins. We performed PCA+IWP analysis on the 200-ns MD trajectories of an FGF2 monomer (126 residues; PDB ID: 1BFG) and HPNAP (144 residues; PDB ID: 1JI4) to derive time scales (*τ*_*w*_) of each protein’s PC modes. Then, we combined the mapped time scales of the ANM modes and the corresponding eigenvalues for these three proteins (ubiquitin, FGF2 and HPNAP) to derive the general power law 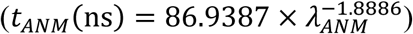 (**Figure S4A**). We would like to acknowledge that the general power law derived here from the three proteins of different sizes (length; also see **Figure S4B**) is nowhere final and presented as a seminal effort for further studies and improvements.

In the **Supplemental Information**, we provide a verification on how the time scales of several dynamics variables can be obtained using aforementioned general power law, as a function of ANM eigenvalue. Thus, a simple molecular timer that characterizes functional motions of biomolecules is obtained. **Figure 4B** summarizes the time scales of discussed experimental observables and ANM modes of interest.

### Derive ANM power law for the sizes of functional motions

Since each ANM mode has been mapped to a PC mode, one can also describe the PC mode variance (the eigenvalue of PC mode) as a function of the corresponding ANM eigenvalue. Combining the aforementioned three protein cases, the linear regression of a log-log plot provides a new power law such that 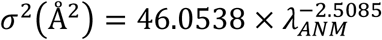 (see the **STAR Methods**). This power law, governing the sizes (variance) of motion as a function of ANM eigenvalues, together with the time power law jointly suggest that the protein’s functional motions follow a *t*^1.33^∼⟨σ^2^⟩ relationship. In the meanwhile, the normal mode theory provides a relation ⟨*σ*^2^⟩ = *k*_*B*_*T*/*mω*^2^ for the harmonic oscillator (where *ω* is the frequency for the normal mode of interest), suggesting that *t*^2^∼⟨*σ*^2^⟩, while the Einstein–Smoluchowski relation (or fluctuation-dissipation theorem) for freely diffused particle in 1D describes a ⟨*σ*^2^⟩ = 2*Dt* relation, or equivalently *t*∼⟨*σ*^2^⟩. Our derived *t*^1.33^∼⟨*σ*^2^⟩ relationship lies closer to free diffusion than to a harmonic oscillator.

### Analysis of ribosomal motions

The power laws are applied to analyze the ribosomal body rotation motion (ratcheting) (Tama et al., 2003; Wang et al., 2004) between the 30S and 50S subunits of the *Thermus thermophilus* ribosome (Tama et al., 2003), which is essential for ribosomal translocation during protein translation (Cornish et al., 2008; Frank and Agrawal, 2000). We perform ANM analysis on the non-rotated ribosome conformation and the 25^th^ slowest ANM mode is identified as the ratcheting motion, see details in the **STAR Methods** and **Supplemental Information**. We deform the ribosome by the size of mode 25 suggested by the variance power law and the rotated ribosome shows a 9° rotation which is close to the experimentally measured rotation of 7° (according to the non-rotated state [PDB ID: 4V6F] and rotated state [PDB ID: 4V9H]) (Jenner et al., 2010; Tourigny et al., 2013).

The estimated timescale of the ribosomal body-rotation (ratcheting) according to simulation (Whitford et al., 2013) and stopped-flow (Cornish et al., 2008; Guo and Noller, 2012) is between >1.3 µs and <5.0 ms, respectively. Individual power laws estimated from ubiquitin and FGF2 predicts the timescale to be between 11.5 µs and 651.9 µs, respectively. However, the HPNAP-derived power law estimates a timescale of less than 1.3 µs, inconsistent with the simulation results (Whitford et al., 2013). Therefore, for predicted time scales longer than 600 ns, which is the longest simulation we conducted to derive the previous power laws, we adopt the power law derived from ubiquitin and FGF2 only, which gives a 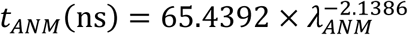 (see **Figure S5**). With this power law, we obtain a timescale of 327.5 µs for the ribosomal ratcheting motion (see timescale estimates for all the slowest ribosomal normal modes in **Table S1**), which is just 1.8-fold longer than 180 µs, the time scale estimated by the power law derived from all the 3 proteins 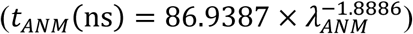. Using the former power law (for estimated time scale > 600 ns), the L1 stalk closing and 30S head swiveling motion are estimated to have a time scale of 202-253 and 128 µs, respectively (see **Supplemental Information)**.

## DISCUSSION

To summarize, the two power laws we derived are combined to give ⟨*σ*^2^⟩ = 0.1223*t*^1.3282^. Such a “guided” (structure-encoded) diffusion motion (*t*∼⟨*σ*^2^⟩^0.75^) is slower than a harmonic oscillator that travels with a (*t*∼⟨*σ*^2^⟩^0.5^) (Atilgan et al., 2001) relation, but faster than the free diffusion (*t*∼⟨*σ*^2^⟩) (McQuarrie, 2000). The predicted time scale using our power law at 100s of microseconds to milliseconds is estimated to have an error of 2-4 fold or less than an order of magnitude. We hope that the introduced method to turn ENMs into a molecular timer and sizer can aid in our quantitative understanding of functional and anharmonic motions of biomolecules. The codes for performing the analysis are available on https://github.com/Hong-Rui/bioStructureM.

## Supporting information

Ubiquitin MD Simulation Input Files

FGF2 MD Simulation Input Files

HPNAP MD Simulation Input Files

## ACKNOWLEDGEMENTS

We acknowledge the computational resources supported by High Performance Computing Infrastructure (HPCI), Japan, by National Center for High-performance Computing (NCHC) of National Applied Research Laboratories (NARLabs) of Taiwan, and by supercomputers at the RCCS, The National Institute of Natural Science, and ISSP, The University of Tokyo. J.C. acknowledges financial supports from MOST and Taiwan International Graduate Program, Academia Sinica, Taipei, Taiwan. This work is funded by the Ministry of Science and Technology (104-2113-M-007-019 and 106-2313-B-007-001-) and National Center for Theoretical Sciences, Taiwan to L.W.Y and by MEXT/JSPS KAKENHI (No. 25104002 and 15H04357) to A.K., as the “Priority Issue on Post-K Computer” (Building Innovative Drug Discovery Infrastructure Through Functional Control of Biomolecular Systems).

## AUTHOR CONTRIBUTIONS

L.W.Y. designed, developed and supervised the project. L.W.Y. and Y.J. implemented the Wiener–Khintchine theorem to obtain time-correlation functions. A.K. and L.W.Y. design the intensity-weighted time period, implemented by H.R.L. with zero paddings. K.T., L.W.Y. and J.C. jointly design the algorithms to compute the order parameters, implemented by H.R.L, K.T. and J.C., who also jointly generate the data for ADPs, RMSFs and (PC)mode-(ANM)mode mapping. K.C.C. performed the ANM analysis for ribosome. Y.Y.C. wrote part of the MATLAB codes for ANM analysis included in the bioStructureM package (https://github.com/Hong-Rui/bioStructureM). H.J.L., J.C. and L.W.Y. jointly drafted the first version of the paper, revised and finalized by all the co-authors.

## STAR METHODS

### CONTACT FOR REAGENT AND RESOURCE SHARING

Further information and requests for resources and reagents should be directed to and will be fulfilled by the Lead Contacts, Lee-Wei Yang or Akio Kitao.

## METHOD DETAILS

### MD Simulations

Detailed simulation protocols have been previously reported (Takemura and Kitao, 2012). Briefly, a 600ns MD simulation for Ubiquitin was performed using the PMEMD module of AMBER 10 (Case et al., 2005) with the ff99SB force field (Hornak et al., 2006), while CHARMM36 (Huang and Mackerell, 2013) force field was used to perform 200ns MD simulations for FGF2 and HPNAP with NAMD 2.10 (Phillips et al., 2005). The TIP3P model was used, and the distance between the outer most protein atom and the closest simulation box face in the initial setup was 20 Å. All systems were brought to thermodynamic equilibrium at 300 K and 1 atm using a weak coupling thermostat and a barostat. The equations of motion were integrated with a time step of 2 fs/step. The long-range Coulomb energy was evaluated using the particle mesh Ewald (PME) method. MD snapshots were stored at a rate no less than one snapshot per 1 picosecond.

### Principal component analysis (PCA) of the covariance matrix computed from MD trajectories

Proteins in the snapshots were iteratively superimposed onto their mean structure, <**R**>, and the covariance matrix, <Δ**R**Δ**R**^***T***^>, is constructed using a deviation matrix **Q** and decomposed into its corresponding eigenvalues and eigenvectors such that:

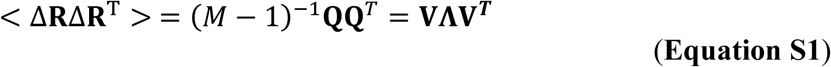

***Δ*R** is the 3*N*-dimensional deviation vector, and **Q** is the 3*N* × *M* matrix, where *N* and *M* are, respectively, the number of protein atoms in the analysis and the number of snapshots. Each column in **Q** represents the deviation of a given snapshot from the mean structure, while each element in that column is the deviation of a given atom in the x-, y- or z-dimension. **V**_3*N*×3*N*_ is the eigenvector matrix containing 3*N* eigenvectors (or principal components, PCs), each of which is 3*N*-dimensional. **Λ** is the 3*N* × 3*N* diagonal matrix of rank-ordered eigenvalues (from large to small).

The snapshots of a trajectory are then projected onto the principal components to form a projection matrix **U**_3*N*×*M*_ as

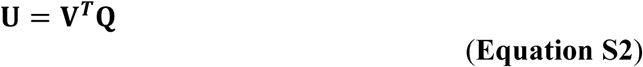

where the row *k* in **U, u**_*k*_ = [*u*_*k,0*_, *u*_*k,1*_, …, *u*_*k,M-1*_], contains the projections of *M* snapshots onto a given PC eigenvector **V**_*k*_ for the *k*’th PC mode. Each snapshot of the protein structure is a scalar value (PC mode coordinate) on the mode *k.*

### Wiener–Khintchine theorem (WKT)

In this section, we show the detailed derivations and implementations of the autocorrelation functions from the PC modes, obtained from MD simulations, using the Wiener–Khintchine theorem. For the *k*^*th*^ mode and a trajectory containing *M* snapshots, we can express the trajectory as a discrete function of time *s* as *u*_*k*_(*s*) = {*u*_*k*0_, *u*_*k*1_, … *u*_*kM*–1_}. Our goal here is to obtain the autocorrelation for the ‘particle motion’ (the projections of the MD snapshots) at the *n*^*th*^ time point. In statistical mechanics, it is, by definition, the ensemble average of the correlation of particle positions at a time shift *t* ≡ *n*Δ*t*, where Δ*t* is the time step. For long-enough equilibrium simulation, the ensemble average can be replaced by the time average under the ergodic hypothesis such that

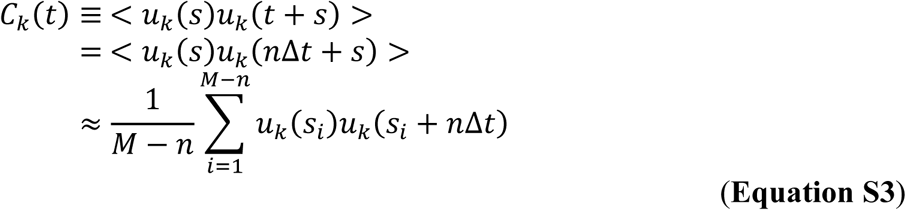

where *s*_*i*_ is the time point when we start to calculate the time-correlation between the *u*_*k*_ at *s*_*i*_ and the *u*_*k*_ after *n* time-steps; *n* goes from 0 to (*M* – 1). The summation runs from 1 to (*M* – *n*) time point since we are only interested in a *n*-time-step shift. The computation time of such a calculation grows with *O*(*M*^*2*^) for all the possible *t* to be obtained. Long computation time is inevitable especially for a long trajectory, repeated for many modes (3*N* – 6, where *N* is the degrees of freedom being analyzed). For our purpose here as well lessening the computational burden, we want to obtain the time-correlation function by first Fourier transforming snapshot projections on a given mode into their counterparts in the frequency domain, and then inverse transform the power spectrum back to obtain the *C*_*k*_(*t*). This mathematical treatment is known as Wiener–Khintchine theorem (McQuarrie, 2000) in statistical mechanics, which proves that the autocorrelation function is equal to the inverse Fourier transformation of the power spectrum of the trajectories such that

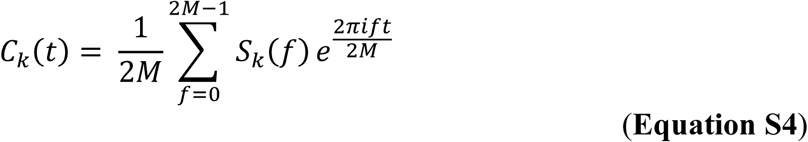

where power spectrum *S*_*k*_(ω) is defined as the magnitude of the Fourier transformation of snapshot projections on the *k*^*th*^ PC mode 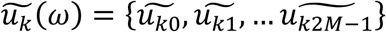, such that 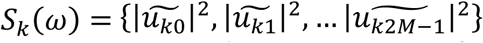. There are 2*M* components in *u*_*k*_(*s*) instead of *M* components after we pad *M* zeros after the original *M u*_*k*_ projections such that *u*_*k*_(*s*) = {*u*_*k*0_, *u*_*k*1_, … *u*_*kM*–1_, 0^1^, 0^2^, … 0^*M*^}. This is to make sure that we obtain *M* positive frequency values. In addition, one can notice that in the last line of **Equation S3**, termed as brute force method herein, for the *n*^*th*^ value of correlation function, the summation goes from 1 to (*M* – *n*), leading to less addends in the summation as *n* increases. Therefore, we need to pad enough zeros at the end of the time series as the input of DFT in order to make sure the Fourier transformation reproduces correct results. In our case, we pad *M* zeros in the end of the sequence so that after the inverse transformation we get back a sequence of length 2*M*. For our needs, we only take the sequence in the positive time domain, with *M* time points. The comparison between Wiener–Khintchine theorem and the brute force method shows that Wiener–Khintchine theorem can indeed reproduce correct time correlation functions, evident that our method is applicable in computing autocorrelation functions with a time complexity of *O*(*M* ln*M*).

### Intensity-Weighted Period (IWP), Relaxation Time and Characteristic Time

The IWP for a specific PC mode *k* is computed as

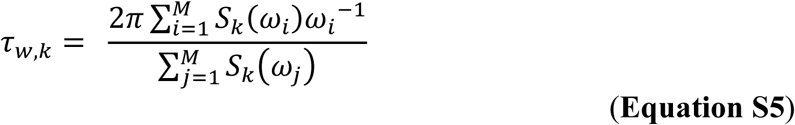

where *S*_*k*_(*ω*_*i*_) is the power for the angular frequency, *ω*_*i*_.

While, the relaxation of *C*_*k*_(*t*) is modeled as an exponential function

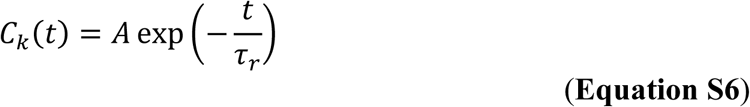

The values of *C*_*k*_*(t)* from unity (when *t* = 0) to the first instance that the function reaches zero (<10^−5^) were used to fit the exponential and obtain *τ*_*r*_.

The characteristic time is defined as

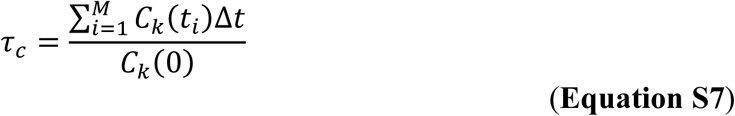

where Δ*t* = 0.1 ps is the time interval between two consecutive snapshots.

### Fluctuation Profile Matching (FPM)

The FPM method is used to identify a subset of PC modes that correspond to dynamical profiles including RMSF, ADP or order parameters. This allows the assignment of timescale and variance of the slowest PC mode to the profile.

FPM takes 2 profiles of the same dynamical variable (one experiment and one theoretical, or two theoretical ones) and assesses their degree of agreement via Pearson’s correlation coefficient (Pearson, 1895), which ranges from −1 to 1, while gradually removing some constituent modes.

FPM is used in the following sections to estimate the timescale of motions producing the RMSF, ADP and order parameter profiles of ubiquitin from the IWP of PC modes. In a later section, FPM is used to assign the timescale and variance to each ANM mode which provides the data for fitting the power laws.

#### RMSF and ADP Profiles for Ubiquitin derived from X-ray/NMR-determined Structures and MD

The RMSF_*exp*_ profile was calculated by treating the ensemble of 32 NMR-determined conformers (PDB ID: 1G6J) of ubiquitin (Cα atoms only) as snapshots and obtaining the <Δ**R**Δ**R**^***T***^>_*exp,1G6J*_ matrix by **Equation S1**. Then the RMSF of each residue is the square-root of the sum of the x-, y- and z-dimension variances for the respective residues taken from the diagonal of <Δ**R**Δ**R**^***T***^>_*exp,1G6J*_. The RMSF_*MD,k*_ profile is computed using the same method but with their respective <Δ**R**Δ**R**^***T***^>_*MD,k*_ matrix in **Equation S8**.

To obtain ADP_*exp*_, the Cα atoms of the ubiquitin X-ray structure PDB ID: 2GBJ (chain B) is superimposed onto the mean structure of MD snapshots. The rotation matrix **R’**_3×3_ used to remove rotational differences between the 2 structures is obtained. Then, the ADP matrix, **U’**_3×3_, for each residue is rotated to obtain the superimposed ADP matrices (**R’U’R’**^***T***^) (Eyal et al., 2007). From the ADP matrix, the ADP dynamical variable of each residue consisting of 3 variance components and 3 unique covariance components (the xy- and yx-components are the same and so are xz- and zx-components, etc.) are extracted to form the 6*N*-dimensional ADP_exp_ profile.

While, the ADP matrix from MD for each Cα atom is the atom’s x-, y- and z-dimension covariance matrix extracted from the full <Δ**R**Δ**R**^**T**^>_*MD,k*_ matrix. The component of this matrix was extracted exactly as above to form the ADP_*MD,k*_ profile.

#### An example of assigning the time-scale of an observed dynamical variable from the IWP of an FPM matched PC mode

For instance, we can take an ensemble of 32 NMR-determined conformers (PDB ID: 1G6J) and form a Cα-only 3*N*×3*N* <Δ**R**Δ**R**^***T***^>_*exp,1G6J*_ matrix to obtain RMSF_*exp*_ (**Figure 3A, top**), see above for details. There are 210 (3*N* – 6 = 210) RMSF_*MD,k*_ profiles, each calculated using all the PC modes ≥ *k*^*th*^ mode. Among the 210 RMSF profiles, RMSF_*MD,k=*25_ has the highest correlation (0.69) with RMSF_*exp*_. Thereby, we claim that the spatial spread of this NMR ensemble takes place within the time scale of *τ*_*w,k=*25_ = 9.32 ns (**Figure 3A, middle and bottom**).

For the case of ADP, ADP_*exp*_ (taken from PDB ID: 2GBJ) has the highest correlation (0.86) with ADP_*MD,k=*41_, see **Figure 3A, top**. PC mode 41 has a *τ*_*w*_ value of 6.17 ns, which we assign as the time scale of the ADP distributions for ubiquitin. The correlation coefficients of RMSF and ADP oFPs compared with the theoretical profiles derived using different *k*’s are shown in **Figure S2A**.

#### Theoretical Covariance Matrix Comprising a set of PC modes Derived from MD simulations

The eigenvectors and eigenvalues are obtained as per **Equation S1** with the first *k*-1 modes (lower PC modes have larger variances or eigenvalues) being removed such that

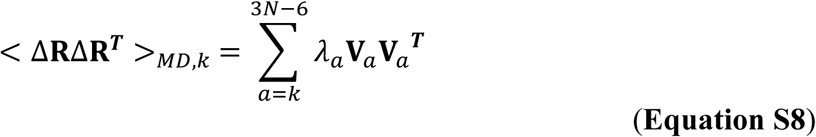

where **V**_*a*_ is the eigenvector with its corresponding eigenvalue, λ_*a*_, taken from the diagonal of **Λ** in **Equation S1** for the *a*’th PC mode.

Take for example, to compute the <Δ**R**Δ**R**^***T***^>_*MD,5*_ covariance matrix, the four PCs with the largest eigenvalues/variances are not included in the summation, thus excluding their contribution to the covariance matrix.

### Order Parameter Profiles derived from NMR-determined structural ensemble and MD snapshots

*S*^2^ describes the order of the backbone -NH bond vector *r*_*ij*_ = (*x*_*ij*_, *y*_*ij*_, *z*_*ij*_) pointing from atom *i* to *j* (herein, *i* is N, and *j* is H) which can be approximated as (Best and Vendruscolo, 2004; Henry and Szabo, 1985; Yang et al., 2009)

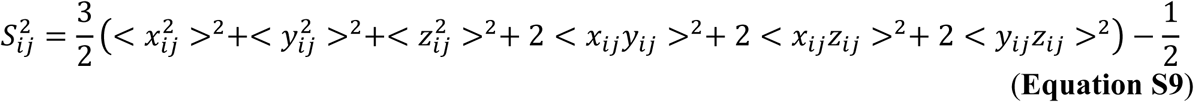

Here, the length of *r*_*ij*_ is normalized to unity. The angular brackets denote averages taken over the *M* snapshots; herein *M* = 6,000,000 or 1,200,000 are for the 600 ns and 120 ns trajectories, respectively.

To obtain the MD-derived *S*^*2*^_*MD,k*_ profile for all the PC modes ≥ *k*^*th*^ mode, the protein structure in each snapshot is rebuilt using all the PC modes ≥ *k*^*th*^ mode which is then used to obtain the -NH bond vectors for calculating *S*^*2*^_*MD,k,ij*_ with **Equation S9**.

The method used to rebuild the structures as a function of constituent modes is as follows, for ubiquitin’s structure in snapshot *m*, the coordinates of its atoms (**R**_*m*_) can be expressed as

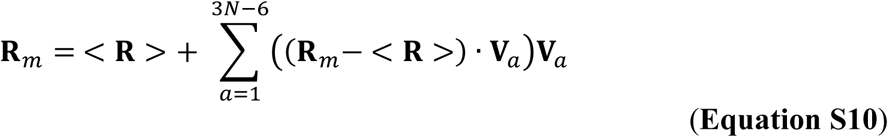

Then, the atom coordinates of snapshot *m* rebuilt using all the PC modes ≥ *k*^*th*^ mode can be obtained by the following equation

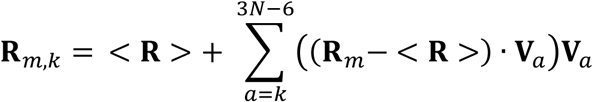

where <**R**> represents the mean coordinate of the structure over *M* snapshots, and **V**_*a*_ is the *a*^*th*^ PC eigenvector.

Now, for each snapshot **R**_*m,k*_, we compute the normalized NH bond vector *r*_*ij*_ for every residue (except for prolines). The averages of *x*_*ij*_^2^, *y*_*ij*_^2^, *z*_*ij*_^2^, *x*_*ij*_*y*_*ij*_, *x*_*ij*_*z*_*ij*_ and *y*_*ij*_*z*_*ij*_ over the whole trajectory were calculated and used to obtain *S*^*2*^_*MD,k,ij*_.

The experimental order parameters, S^2^_*exp*_, for ubiquitin were taken from Tjandra et al (Tjandra et al., 1995).

### Derivation of General Elastic Network Model (ENM) – the relation between covariance and Hessian

In the equilibrium state, the protein system is harmonically approximated and assumed to have the Hamiltonian

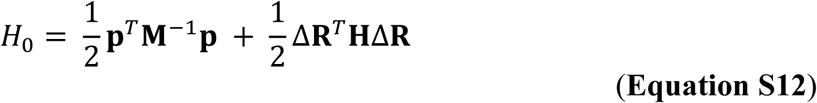

Where **p** is the 3*N*×1 momentum vector in the Cartesian coordinate system, and **M**^−1^ is the inverse of a 3*N*×3*N* diagonal mass matrix comprising elements of triplicate mass of every node (Yang, 2011; Yang and Chng, 2008; Yang et al., 2014). The superscript *T* denotes matrix transposition. Δ**R** is a 3*N*×1 displacement vector and **H** is the symmetric Hessian matrix, or, the force constant tensor of the system, and have the meaning of coupling force constants in each pairwise connected degrees of freedom.

Here we note that since **H** is diagonalizable, for the ease of the following derivations of the ensemble average of positional covariance, we transform **H** in normal-mode space, such that

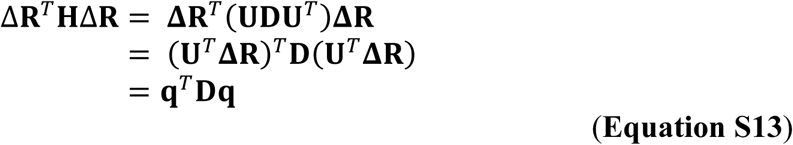

Where **D** is the diagonal eigenvalue matrix, **U** is the unitary eigenvector matrix and **q** is the displacement vector in normal-mode space with the relationship **q** = **U**^*T*^**ΔR**.

Since unitary transformation conserves length so that we can change the integration to normal-mode space, which is readily solvable.

Now, from **eqs S12 and S13**, we write *H*_0_ in the expanded form as

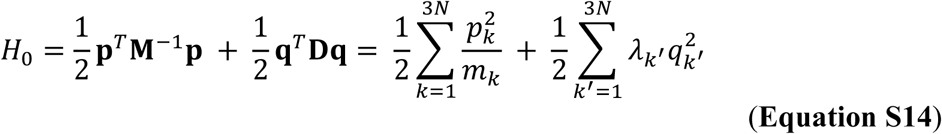

In the above expression, *k* and *k’* represent the index through all 3*N* degrees of freedom, and *m*_*k*_ is the mass elements in **M** with *N* triplicates.

In canonical ensemble we write down the partition function with phase space formulation and solve it by expanding all the terms

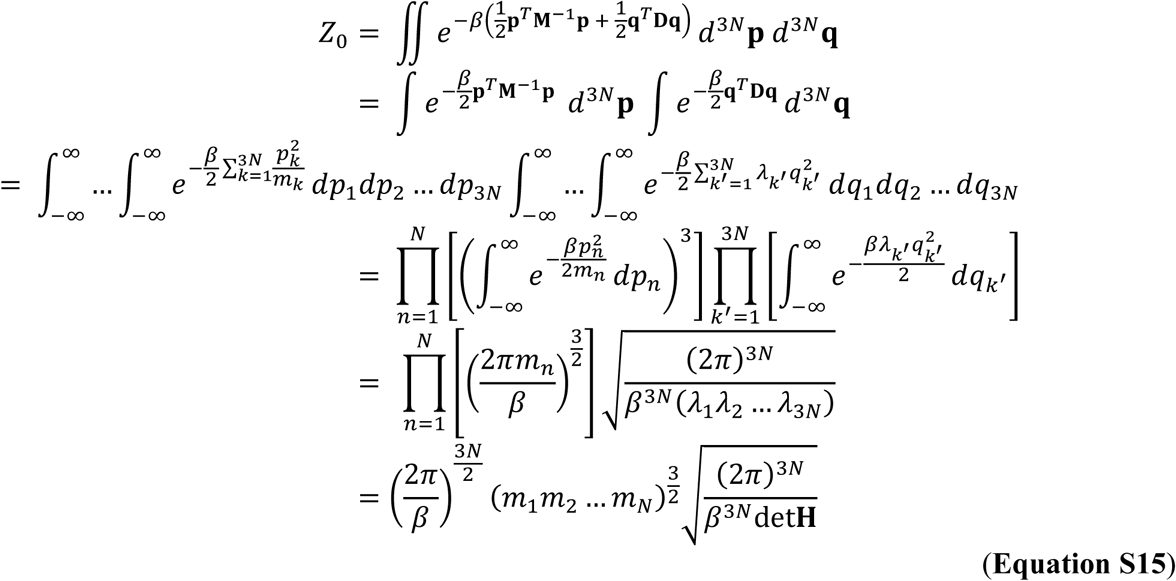

where β=1/*k*_*B*_*T*; *k*_*B*_ is the Boltzmann constant and *T* is the absolute temperature.

In the equation above we use the property that for a symmetric matrix the product of its eigenvalues is the determinant of itself.

To calculate the positional covariance, we note one derivative

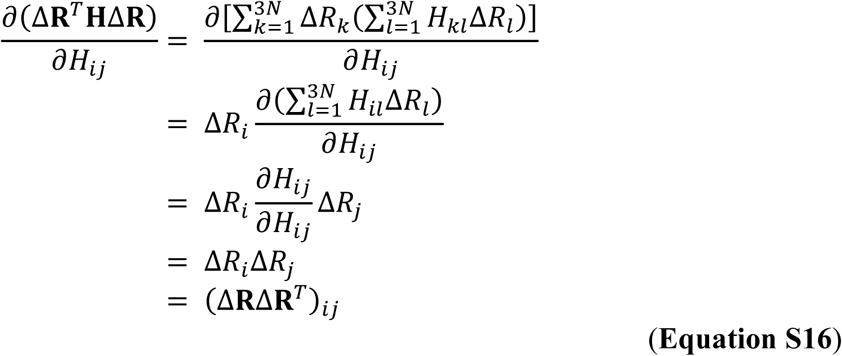

Now

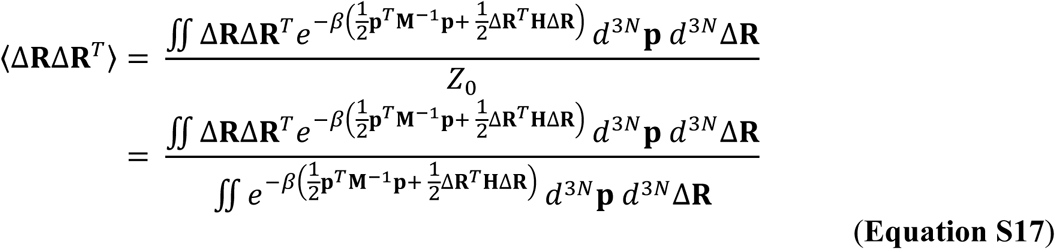

For each element in the covariance matrix

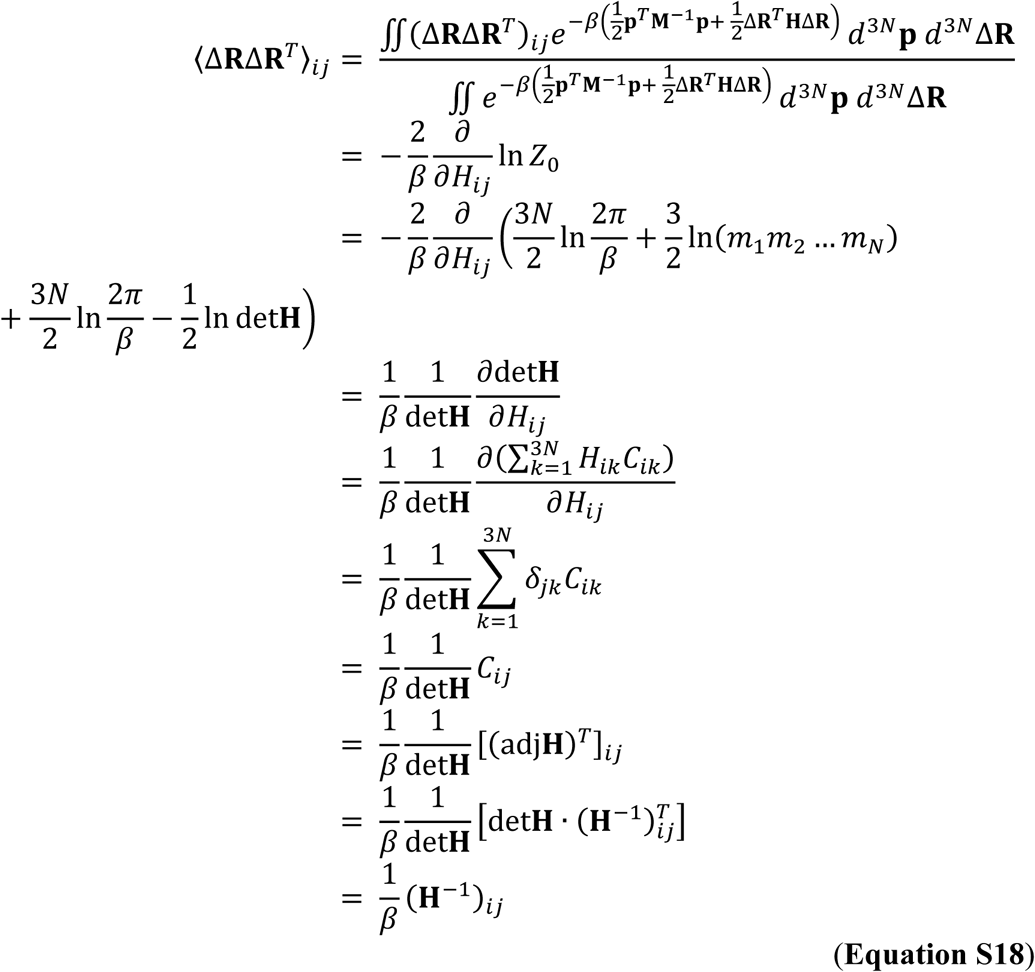

Where *C*_*ik*_ in the above derivation is the *ik*-element of the cofactor matrix of the Hessian. Here we use Laplace formula to expand the Hessian determinant and using the properties that (1) the adjugate matrix is the transpose of cofactor matrix, (2) inverse of any nonsingular matrix **A** is 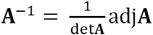 adj**A** and (3) the inverse of any symmetric matrix, in our case Hessian, is still symmetric.

Finally, in matrix form we write

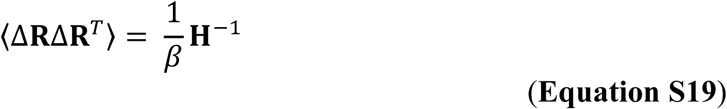

which is the expected result by analogy to one-dimensional harmonic oscillator system.

This result holds true for all types of ENMs as long as the system is harmonically approximated. For ENMs that have translational and/or rotational invariance (Yang, 2011), which makes the rank of **H** less than its dimension, **H**^−1^ is solved by eigenvalue decomposition followed by removing the terms containing zero eigenvalues.

### The Anisotropic Network Model (ANM)

The Anisotropic Network Model (ANM) is a type of ENM with the following potential energy function

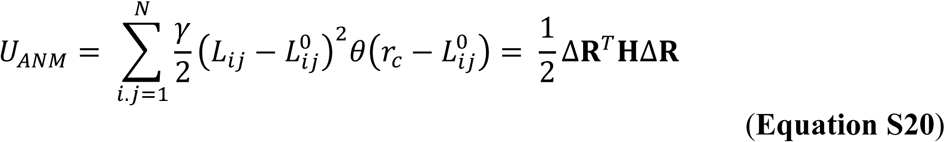

where the index *i* and *j* represents the nodes (Cα atoms) that runs from 1 to *N*. The γ represents the universal spring constant of the system. *L*_*ij*_ is the current distance between the *i*^*th*^ and *j*^*th*^ nodes while *L*_*ij*_^0^ is the equilibrium distance between *i* and *j*. When *L*_*ij*_^0^ is ≤ *r*_*c*_, the cut-off distance, then these two nodes are treated as connected. This is expressed as a Heaviside-step function *θ* in **Equation S20**.

All the elements of the Hessian matrix **H** can be calculated as the second derivative of the potential energy function. Details of this can be found in the supplementary information of our previous work (Yang, 2011). The eigenvector (***V***_*ANM*_) and eigenvalue (*λ*_*ANM*_) of each ANM mode is calculated from the eigen-decomposition of **H such that**

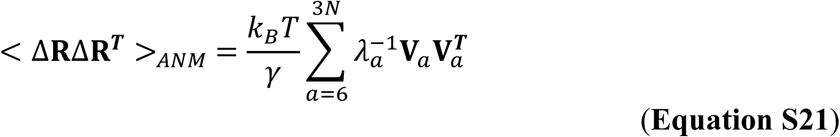

ANM analysis can be performed on the DynOmics Portal 1.0 (Li et al., 2017): http://dyn.life.nthu.edu.tw/oENM/.

### Theoretical Covariance Matrix, RMSF and ADP from ANM With Removed Modes

The ANM derived covariance matrix for all ANM modes ≥ *l* (lower ANM modes have lower frequencies or smaller eigenvalues) can be calculated as follows

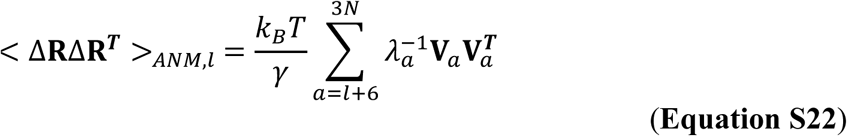

where ***V***_*a*_ and *λ*_*a*_ are the eigenvector and eigenvalue of the *a*’th ANM mode.

The RMSF_*ANM,l*_ and ADP_*ANM,l*_ profiles were computed from the <Δ**R**Δ**R**^***T***^>_*ANM,l*_ matrix with the same methods used for MD-derived profiles of RMSF and ADP.

### ANM Mode Time Scale Assignment Through ANM-PCA Mode Mapping

Every ANM mode *l* was mapped to a PC mode *k\** from MD by comparing the RMSF_*ANM,l*_ with the RMSF_*MD,k*_ profiles for different *k*’s using FPM to identify the *k* = *k\** with the highest correlation. Every ANM mode *l* was then assigned the IWP (*τ*_*w,k\**_) of the mapped PC mode *k\**.

The ANM-PCA mode mapping might result in a PC mode getting mapped to multiple ANM modes. The ANM mode with the highest correlation with the PC mode was kept while the rest were removed. This filtered set of ANM-PCA mode mapping was used to fit the power laws.

### Fitting the Time Power Laws

The relationship between an ANM mode’s eigenvalue (*λ*_*ANM*_) and mapped time scale (*t*_*ANM*_) was modeled as a power law in the form of *t*_*ANM*_ = *c* × *λ*_*ANM*_^*d*^, with the constants *c* and *d* to be found through the fitting procedure described below.

The natural log of the power law, ln (*t*_*ANM*_) = ln (*c*) + *d* ln (*λ*_*ANM*_), was used to get a linear function of ln (*t*_*ANM*_) with respect to ln (*λ*_*ANM*_). Least squares fitting was then used to obtain the parameters *c* and *d*.

### Fitting the Variance Power Law

Given the mapping between ANM modes and PCA modes, the *λ*_*ANM*_ and *λ*_*PCA*_ (PC mode’s eigenvalue/variance) were used to fit a variance power law such that *λ*_*PCA*_ = *a* × *λ*_*ANM*_^*b*^, where *a* and *b* are constants. The constants were found by following the same procedure used to fit the time power laws.

### Identifying the Functional Modes Corresponding to Ribosomal Body Rotation (Ratcheting) and Head Swiveling Motions

The body rotation is known to be generally around an axis that connects the mass centers of the 30S and 50S subunits of the ribosome (Kurkcuoglu et al., 2008; Tama et al., 2003; Wang et al., 2004). Thus, the axis of rotation, **r**_*B*_ the vector connecting the COMs of the 2 subunits as observed in the non-rotated x-ray crystallographically resolved conformation (PDB ID: 4V6F). While for the head swiveling motion’s axis of rotation, **r**_*H*_, it is the vector between the COMs of the 30S body and 30S head. Where the 30S head is defined as the residues 921-1396 of the 16S rRNA complexed with the ribosomal proteins S3, S7, S9, S10, S13, S14, S19 and Thx. While, the 30S body is defined as residues 5-920 and 1399-1543 of the 16S rRNA complexed with ribosomal proteins S2, S4, S5, S6, S8, S11, S12, S15, S16, S17, S18 and S20.

ANM analysis was performed on the non-rotated x-ray solved conformation. The *k*^th^ ANM eigenvector (**V**_*k*_) was used to deform the non-rotated conformation by

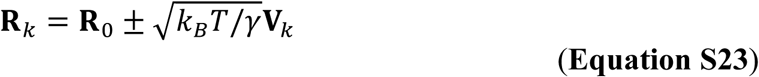

where **R**_*0*_ is a 3*N* vector containing the coordinates of the *N* atoms in the non-rotated conformation and **R**_*k*_ is the conformation deformed from **R**_*0*_ along the *k*^th^ ANM mode by setting *k*_*B*_*T*/γ to be unity. For the purpose of obtaining axis of rotation, the results do not change as long as *k*_*B*_*T*/γ is set to be a small value.

The ratcheting motion is relative to the 50s subunit and therefore the deformed conformation was first superimposed at the 50S subunit. To capture the head swiveling motion, we superimposed the deformed conformation at the 50S subunit and 30S’ body part to avoid capturing the ratcheting motion and examined the rotation of the 30S head.

Considering the transition from non-rotated (**R**_**0**_) to deformed conformation (**R**_**k**_) takes place at a time interval *Δt* (in seconds), the angular momentum vector (**L**_*k*_) for the *k*^th^ mode is the cross product between the mass-weighted position of atoms and atom velocity, such that

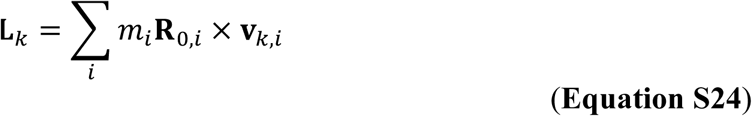

where **R**_*0,i*_ is the *i*^th^ atom’s position vector in the non-rotated conformation, *m*_*i*_ is the atom mass of *i* and the velocity vector **v**_*k,i*_ = (**R**_*k,i*_ – **R**_*0,i*_)/*Δt* where **R**_*k,i*_ is the position vector of atom *i* in the conformation deformed along the *k*^th^ mode.

**L**_*k*_ is related to the angular velocity vector (**ω**_*k*_) by

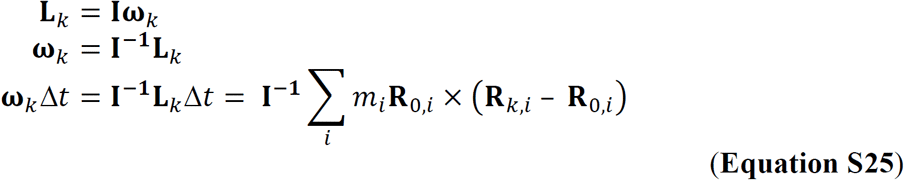

The direction of **ω**_*k*_ is the *axis of rotation* for mode *k* while |**ω**_*k*_| is the angular velocity in the unit of radians/second and therefore |**ω**_*k*_|*Δt* gives us the angle of rotation in radians (the length of *Δt* does not change the result).

The deviation angle *θ* between **r**_*B or H*_ and rotation axis **ω**_*k*_ from mode *k* were computed for the slowest 50 ANM modes, where cos *θ* = (**r**_*B* *or* *H*_ · **ω**_*k*_)/(||**r**_*B* *or* *H*_|| ||**ω**_*k*_||). The mode with the smallest angle deviating from **r**_*B*_ (or **r**_*H*_) was chosen as the functional mode (*k*^*f*^) for the ratcheting (or for swiveling) motion.

Among the slowest 50 ANM modes, the *rotation axis* derived from mode 25 was identified to have the smallest deviation angle (*θ* = 11.6°) from the vector that connects the mass centers of 30S and 50S subunits. Therefore, mode 25 represents the ribosomal ratcheting motion. As for the head swiveling motion, ANM mode 32 has the smallest angle (*θ* = 3.9°) deviating from the axis of rotation.

### Predicting the Conformation of the Ribosome After the Ratcheting and Head Swiveling Motions

After identifying the ANM mode (*k*^*f*^) that corresponds to the body rotation/head swiveling motion, the conformation of the ribosome could be obtained by

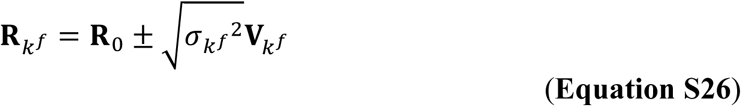

where variance 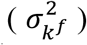 of mode *k*^*f*^ can be predicted by the variance power law 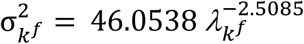 with 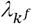, the eigenvalue of mode *k*^*f*^.

### Identifying the Functional Modes and Conformations Corresponding to the L1 Stalk Closing Motion

The L1 stalk is defined as the helices 76-78 (residues 2093-2196) of the 23S rRNA complexed with L1 protein and the COM of E-site is defined by the COM of E-site tRNA as observed in the non-rotated ribosome (PDB ID: 4V6F) (Fei et al., 2008).

To identify the L1 stalk’s functional mode and conformation, the non-rotated ribosome (PDB ID: 4V6F) was first deformed along all 50 modes using the variance power law as in **Equation S26**. The deformed conformations were then superimposed at the 50S subunit as described above. Finally, we identify the mode with the shortest distance between the COM of the deformed L1 stalk and the E-site in the non-rotated ribosome.

The conformations deformed with ANM mode 27 and 28 have the two closest distance (∼26 Å and ∼30 Å, respectively) between the COM of L1 stalk and the COM of the E-site in the ribosome. We treat both modes as part of the L1 stalk closing motion.

## QUANTIFICATION AND STATISTICAL ANALYSIS

To evaluate the quality of the fit between the power laws and timescale/variance, the Pearson correlation coefficient (Pearson, 1895) between the predicted timescale/variance with the MD derived timescale/variance was compared.

The Person correlation coefficient is also used in FPM to evaluate the goodness of fit between two profiles of dynamics variables.

## DATA AND SOFTWARE AVAILABILITY

The input files used to perform the MD simulations are included as zipped files as part of the Supplementary Information.

ANM, WKT, IWP, FPM and the power laws are implemented in Matlab and the codes are available at https://github.com/Hong-Rui/bioStructureM. The MD trajectories of Ubiquitin, FGF2 and HPNAP can be requested by e-mail.

## KEY RESOURCES TABLE

| REAGENT or RESOURCE | SOURCE | IDENTIFIER |
| --- | --- | --- |
| Software and Algorithms |  |  |
| Matlab R2017a | MathWorks | https://www.mathworks.com/ |
| AMBER 10 | (Case et al., 2005) | http://ambermd.org/ |
| NAMD 2.10 | (Phillips et al., 2005) | http://www.ks.uiuc.edu/Research/namd/ |
| Wiener–Khinchine theorem (WKT) | (McQuarrie, 2000) | https://github.com/Hong-Rui/bioStructureM |
| Intensity Weighted Period (IWP) | This paper | https://github.com/Hong-Rui/bioStructureM |
| Fluctuation Profile Matching (FPM) | This paper | https://github.com/Hong-Rui/bioStructureM |
| Power Laws | This paper | https://github.com/Hong-Rui/bioStructureM |

