## Supplemental Information for "An efficient timer and sizer of biomacromolecular motions"

### Supplemental Figures and Table

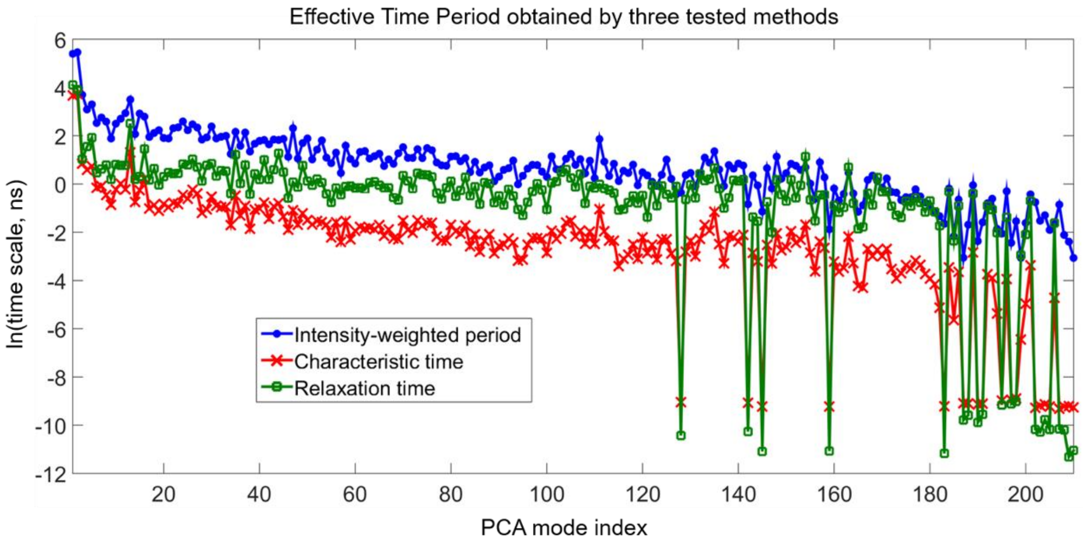


**Figure S1. Time scale of each PC mode of ubiquitin estimated using the three tested methods.** There are 3 × 72 – 6 = 210 PC modes ($C\alpha$-based) calculated from the 600-nanosecond simulation of ubiquitin. The blue, red and green lines show intensity-weighted period ($\tau_w$), characteristic time ($\tau_c$) and relaxation time ($\tau_r$) respectively for all the PC modes.

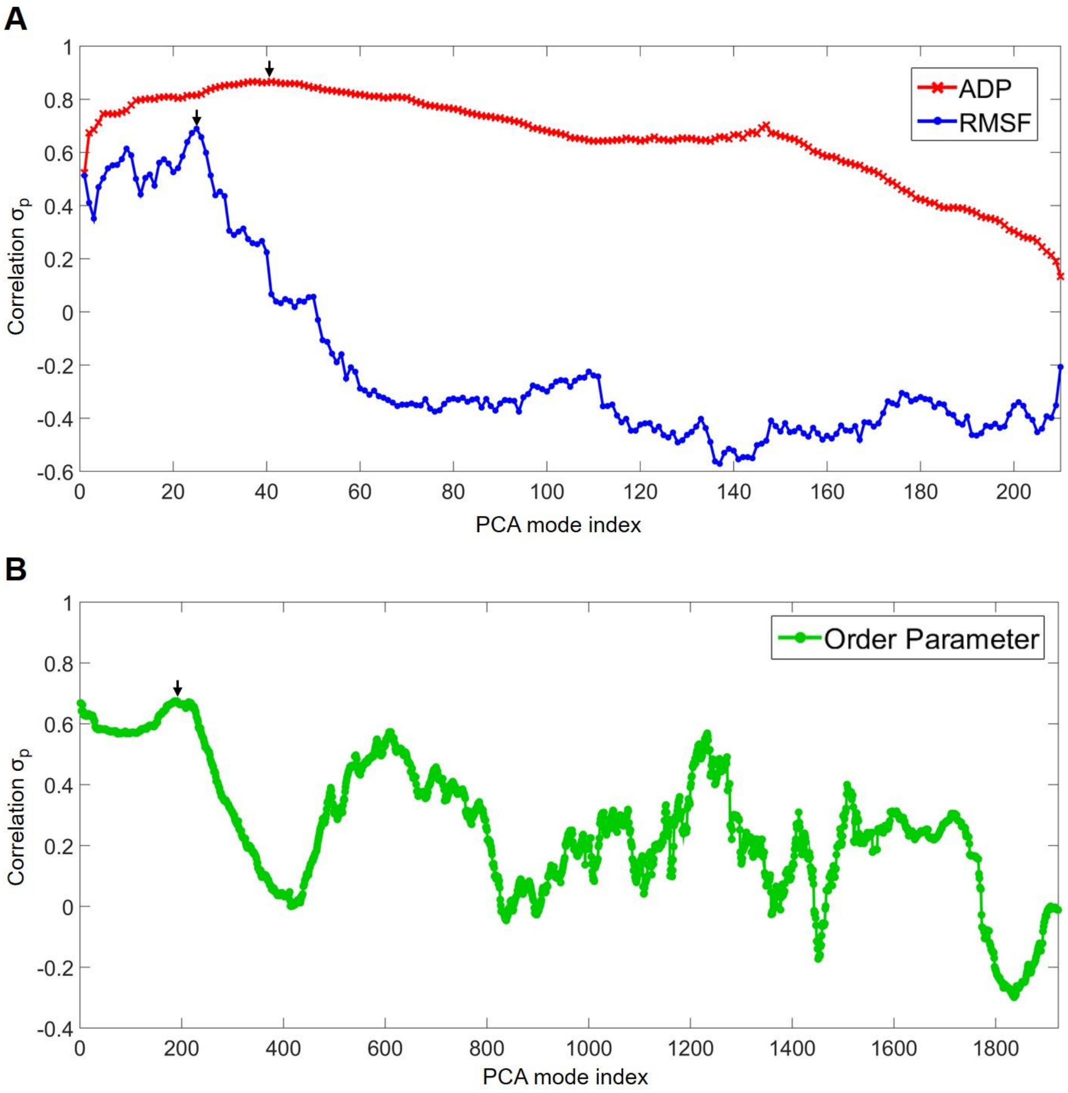


**Figure S2. The PC modes that best match the dynamics variables of ubiquitin.**
**(A)** The Pearson correlation coefficient between the anisotropic displacement parameters (ADP) profile assembled from all the PC modes higher or equal to a specific mode with the experimental ADP profile (PDB: 2GBJ) is shown in red crosses. Blue circles give correlation coefficients between RMSF profile derived from covariance comprising PC modes higher or equal to a given mode (**Equation S8**) and RMSF derived from a set of NMR-determined conformers (PDB: 1G6J). In **(A)**, the PC modes are calculated using only C$\alpha$ atoms (coarse-grained). **(B)** Pearson correlation coefficient between experimental NMR order parameters and order parameters estimated from the PCA of heavy-atoms with the backbone nitrogen's hydrogen atom. The black arrow indicates the PC mode with the highest correlation.

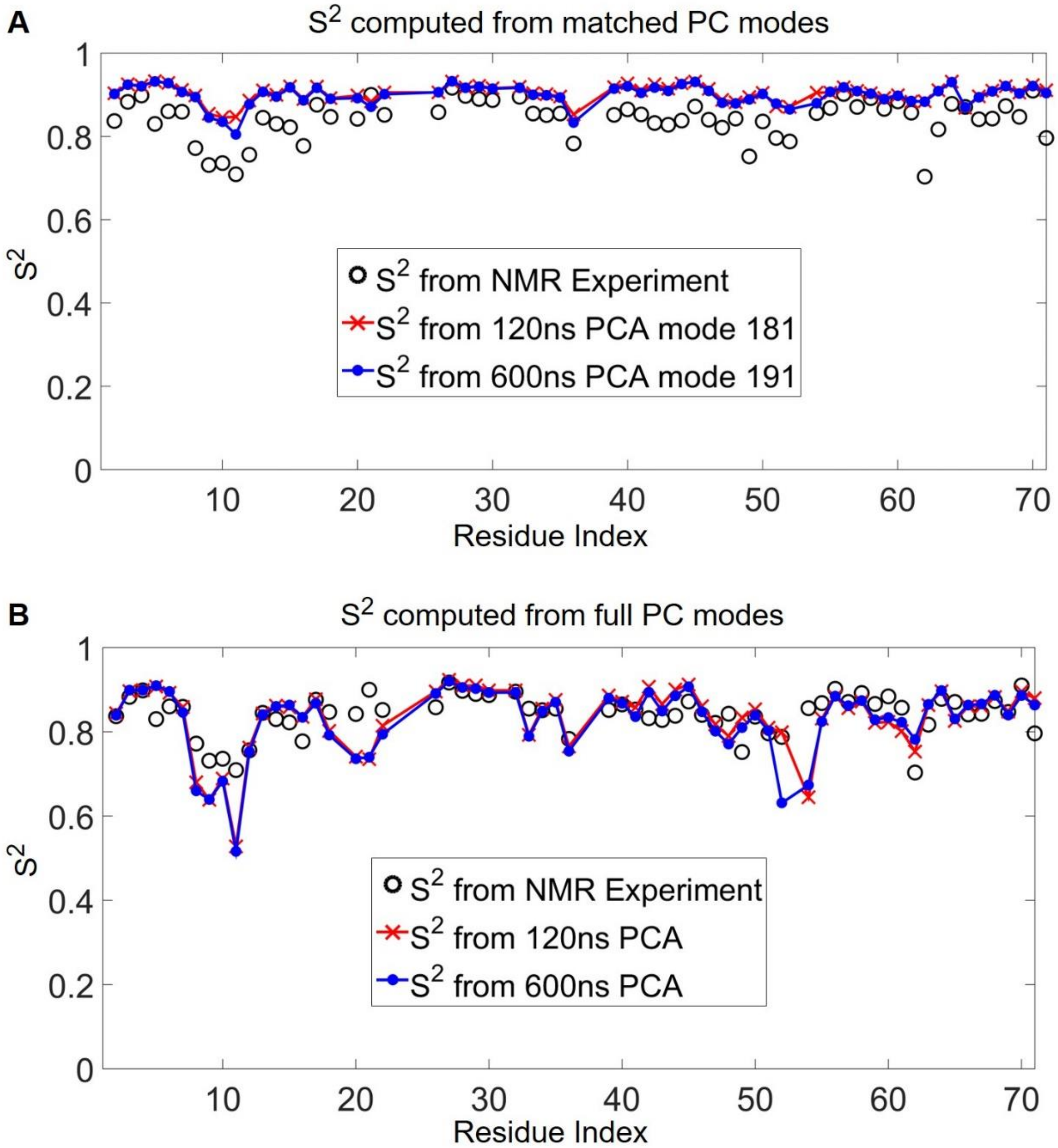


**Figure S3. NMR order parameters ($S^2$) estimated from the 120 ns and 600 ns MD trajectories.**
**(A)** The profiles are the order parameters derived from PC modes ($k$=181 and 191 for 120 and 600ns simulations, respectively) that best-match the experimental data. One can obtain the time scales of 1.0 ns and 2.3 ns for the NMR-characterized $S^2$ profile from the best-matched mode 181 and 191, respectively. **(B)** The profiles are order parameters derived from full PC modes ($k$=1 in **Equation S9**). $S^2_{MD,k=1}$ shows slightly larger (less ordered) bond fluctuations than $S^2_{exp}$ (**Figure S3b)**, which implies that within the 600 and 120-ns timeframes, the protein samples a wider conformational space than that sampled during the NMR relaxation experiment. It is readily understandable that the longer the simulation is, the wider the spatial distribution (and therefore less order) of an atom would result.

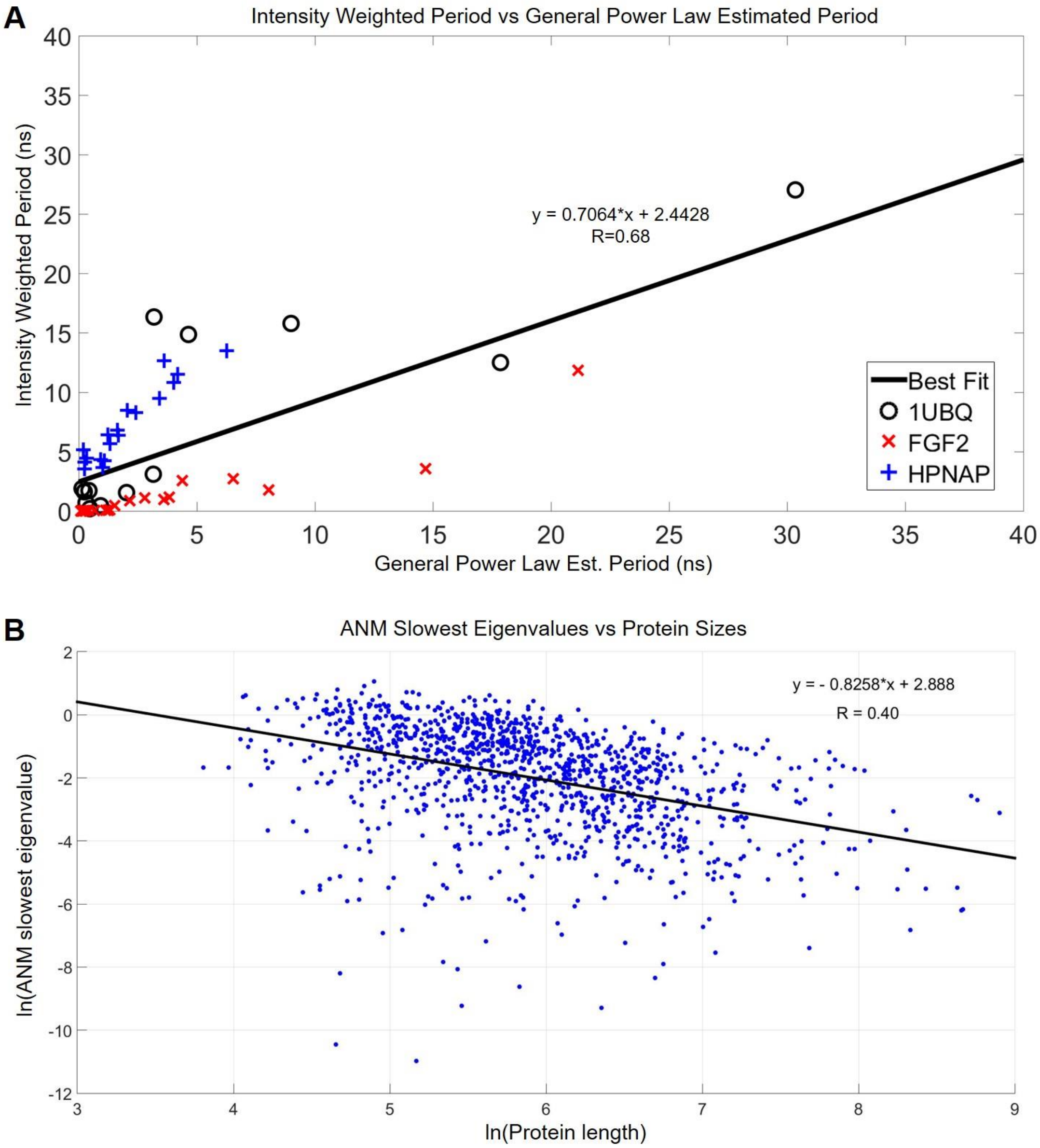


**Figure S4. Comparing IWPs to power law estimated periods and plotting eigenvalues of ANM's slowest modes to the lengths of proteins.**

**(A)** The IWPs are compared to the power-law estimated periods for the combined ANM modes of ubiquitin, FGF2 and HPNAP ($\sigma_p$ = 0.68). **(B)** The ANM eigenvalues of the slowest modes are plotted against protein sizes (the number of residues in a protein) for a previously published set of non-homologous 1228 proteins (Yang et al., 2006), suggesting a moderate correlation; hence the derived power laws as a function of eigenvalues implicitly take account of the size factors of proteins.

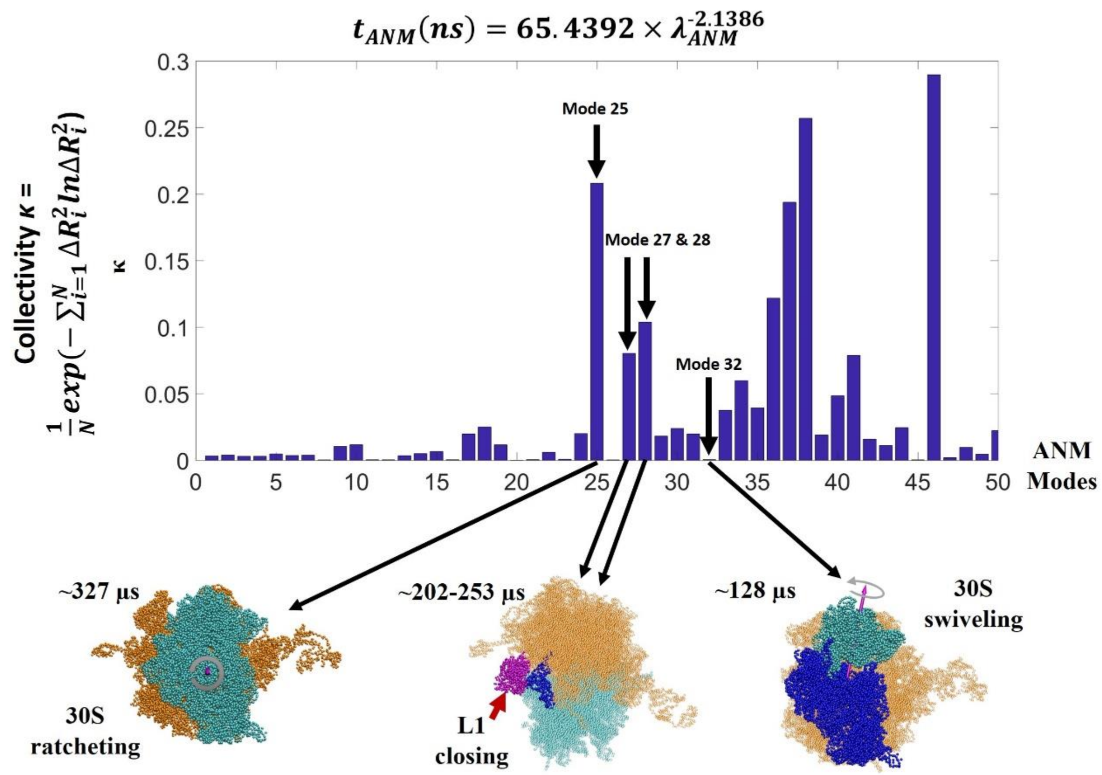


**Figure S5. The collectivity (κ) of the top 50 slowest ANM modes where four of which are the discussed functional motions in ribosome.**

The collectivity (κ) (Tama and Sanejouand, 2001) quantifies the amount of global motion in each ANM mode. The 30S ratcheting motion (mode 25) and the L1 stalk closing motion (modes 27 and 28) are found to be highly collective but not the local 30S head swiveling (mode 32). Only ubiquitin and FGF2 are used to fit the time power law ( $t_{ANM}(\mathrm{ns}) = 65.4392 \times \lambda_{ANM}^{-2.1386}$) for estimating the timescales of these ribosomal motions. 50S of the ribosome is in reddish color.

**Table S1. The top 50 slowest ribosomal motions characterized by ANM**

| | ANM Mode Index | Eigenvalues | Estimated Time Scale (μs) |
|---|---|---|---|
| Slowest motion | 1 | 4.46E-05 | 131624834.27 |
| | 2 | 6.50E-05 | 58969480.25 |
| | 3 | 8.38E-05 | 34246012.91 |
| | 4 | 1.17E-04 | 16796154.77 |
| | 5 | 3.25E-04 | 1882206.56 |
| | 6 | 4.08E-04 | 1161959.99 |
| | 7 | 5.57E-04 | 596328.57 |
| | 8 | 7.64E-04 | 303487.58 |
| | 9 | 8.93E-04 | 217351.77 |
| | 10 | 1.21E-03 | 112995.01 |
| | 11 | 1.50E-03 | 71215.70 |
| | 12 | 3.69E-03 | 10441.04 |
| | 13 | 3.94E-03 | 9065.73 |
| | 14 | 4.07E-03 | 8462.59 |
| | 15 | 5.35E-03 | 4722.31 |
| | 16 | 6.34E-03 | 3288.69 |
| | 17 | 7.45E-03 | 2323.78 |
| | 18 | 8.56E-03 | 1729.35 |
| | 19 | 9.20E-03 | 1480.40 |
| | 20 | 1.10E-02 | 1006.53 |
| | 21 | 1.12E-02 | 974.11 |
| | 22 | 1.22E-02 | 812.59 |
| | 23 | 1.62E-02 | 439.80 |
| | 24 | 1.73E-02 | 385.09 |
| Ratcheting motion | 25 | 1.86E-02 | 327.46 |
| | 26 | 1.91E-02 | 310.47 |
| L1 stalk motion | 27 | 2.10E-02 | 253.48 |
| L1 stalk motion | 28 | 2.33E-02 | 202.40 |
| | 29 | 2.54E-02 | 168.47 |
| | 30 | 2.62E-02 | 158.06 |
| | 31 | 2.76E-02 | 141.51 |
| Head swiveling motion | 32 | 2.90E-02 | 127.57 |
| | 33 | 3.15E-02 | 106.21 |
| | 34 | 3.51E-02 | 84.65 |
| | 35 | 3.81E-02 | 71.06 |
| | 36 | 4.17E-02 | 58.48 |
| | 37 | 4.31E-02 | 54.58 |
| | 38 | 5.26E-02 | 35.59 |
| | 39 | 5.32E-02 | 34.74 |

| | | |
|---|---|---|
| 40 | 5.56E-02 | 31.56 |
| 41 | 5.69E-02 | 30.08 |
| 42 | 6.03E-02 | 26.58 |
| 43 | 6.48E-02 | 22.79 |
| 44 | 6.56E-02 | 22.16 |
| 45 | 6.80E-02 | 20.52 |
| 46 | 6.91E-02 | 19.82 |
| 47 | 7.32E-02 | 17.57 |
| 48 | 7.37E-02 | 17.29 |
| 49 | 7.41E-02 | 17.08 |
| 50 | 7.47E-02 | 16.81 |

ANM is performed on the *Thermus thermophilus* 70S ribosome whose missing atoms, residues and subunits were patched using the individually solved subunits as templates (Chang et al., 2015). The estimated time scales of the vibrational modes are obtained from the time power law $t_{ANM}(\text{ns}) = 65.4392 \times \lambda_{ANM}^{-2.1386}$. See **Figure S5**.

### Supplemental Results

#### Verification of the General ANM Power Law via the Dynamics Analysis of Ubiquitin, FGF2 and HPNAP

To validate the applicability of the general power law, $t_{ANM}(\text{ns}) = 86.9387 \times \lambda_{ANM}^{-1.8886}$, we examined the estimated time scales of the experimental RMSF and ADP profiles (**Figures 3A and 3B**) using ANM-based power law instead of IWP derived from the long MD trajectories. The results showed that theoretical ADPs predicted from removing the first 7 ANM modes agree the best with the experimental ADPs. By substituting the eigenvalue of mode 8 in the general power law, we obtain a time scale of 5.38 ns, which is close to the earlier MD/IWP result of 6.17 ns. In a similar vein, the RMSF profile derived from the 32 NMR conformers (PDB ID: 1G6J) matches the best with the theoretical RMSF comprising the ANM mode 4 and above, which maps to a time scale of 8.70 ns according to the general power law. This is close to the earlier MD/IWP result of 9.32 ns (mapped to the $25^{th}$ PC mode). Furthermore, for FGF2 and HPNAP, we found the time scale of each ANM mode predicted by the general power law closely agrees with that estimated from FPM+WKT, with correlations of 0.95 for FGF2 and 0.93 for HPNAP, respectively.

#### The Size, Conformation and Time Scale of Ribosomal Motions Predicted with the Variance and Time Power Laws

*Finding the relevant modes that describe 30S head swiveling and L1 stalk closing motion of the ribosome and characterizing the size and time scales of these motions*

30S head swiveling motion and L1 stalk motion are also known to be involved in the ribosomal translocation process (Frank and Agrawal, 2000). The 30S head is defined as the 3' major domain of the 16S rRNA spanning residues 921-1396 and the complexed proteins (Mohan et al., 2014) (colored in cyan at the bottom right of **Figure S5**). Its rotational motion relative to the body domain, coined as the swiveling motion, can swivel up to 18° after binding EF-G and the concomitant GTP hydrolysis. This intra-subunit motion together with the inter-subunit ratcheting of the whole 30S drives tRNA translocation (Ratje et al., 2010). Consequently, when the swiveling motion is impeded by an obstacle, such as an mRNA pseudoknot, the dissociation of E-site tRNA and EF-G can be hindered (Caliskan et al., 2014).

On the other hand, L1 stalk is defined as the complex between the helices 76-78 of 23S rRNA and the L1 ribosomal protein (Fei et al., 2008). The L1 stalk motion is not only influenced by the 30S motions (Fei et al., 2008) and the downstream mRNA structures (Chen et al., 2013) but also by its binding to the P-site tRNA (L1 stalk closing) (Trabuco et al., 2010) and guiding it from the P-site to the E-site which involves motions as large as ~20 Å (Valle et al., 2003).

By applying the power laws, we are not only interested in reproducing observed sizes of conformational changes but also provided estimated timescales for these intrinsic and functional motions in ribosome.

A similar method used to identify the ANM mode that corresponds to the ratcheting motion was applied to find the corresponding mode for the swiveling motion (**STAR Methods**). The main differences are that the axis of rotation for head swiveling is defined as the vector pointing from the COM of the 30S body to the COM of the 30S head and the deformed conformer is superimposed at the 50S and 30S body of the non-rotated ribosome (PDB ID: 4V6F), see **STAR Methods** for details. As mentioned in the previous section, we can calculate the rotation axis for each of the 50 slowest ANM modes. The axis of rotation computed for ANM mode 32 has the smallest angle ($\theta$ = 3.9°) deviating from the axis of rotation of the swiveling motion. Deforming the conformation along ANM mode 32 scaled by the variance power law results in an angle of rotation of 3.2° in contrast to the observed 4.4° rotation in the corresponding structures (PDB ID: 4V6F and PDB ID: 4V9H). The predicted time scale using the eigenvalue of mode 32 and the time power law is ~128 μs.

The L1 stalk interacts with the tRNA at the E-site of the ribosome during the ratcheting and the swiveling motions. To identify the ANM mode(s) corresponding to this motion, each of the top 50 slowest ANM modes was scaled by the variance power law and used to deform the non-rotated conformation (PDB ID: 4V6F), see **STAR Methods** for details. The conformations deformed with ANM mode 27 and 28 have the two closest distance (~26 Å and ~30 Å, respectively) between the COM of L1 stalk and the COM of the E-site in the ribosome with the predicted time scales of ~202-253 μs. The obtained distance of ~26-30 Å is closer to the rotated conformation (25 Å; PDB ID: 4V9H) while the corresponding distance in the non-rotated ribosome (PDB ID: 4V6F) is 42 Å.

The fact that the timescale of the ratcheting motion (~327 μs) is closer to the L1 stalk closing (~202-253 μs) than the swiveling motion (~128 μs) suggests a stronger coupling between the ratcheting and the L1 stalk closing, whereas the 30S head moves more independently from the other two. Indeed, the correlation analysis over a dozen of cryo-EM and x-ray solved structures carried out by Agirrezabala et al. (Agirrezabala et al., 2012) showed that the correlation between the ratcheting angles and the swiveling angles is about 0.19 (*p-value* = 0.4241) while that between ratcheting and L1 stalk closing is 0.99 (p-value < 0.0001)

*Collectivity of the ribosomal motions*

Tama and Sanejouand's definition of collectivity (how global a motion is) was applied to each of the ANM modes (Tama and Sanejouand, 2001). As expected, global ratcheting and L1 stalk closing (which is coupled to ratcheting) motions observed in ANM modes 25, 27 and 28 are among the modes with the highest collectivity within the slowest 50 ANM modes (**Figure S5**), whereas the independent local motion of head swiveling has a relatively low collectivity.
